# Combined MEG and EEG suggest a limbic source network of the P3 including retrosplenial cortex and hippocampus

**DOI:** 10.1101/2025.03.24.645008

**Authors:** Diptyajit Das, Matti S. Hämäläinen, Andrew R. Dykstra, Andre Rupp, Alexander Gutschalk

**Author notes:** Corresponding author: E-mail address (A. Gutschalk).

## Abstract

The event-related P3 is evoked by most task-relevant or salient stimuli, but its neural sources remain controversial. Here we reevaluate the role of hippocampus, unambiguously demonstrated in intracranial recordings, and retro-splenial cortex as sources of the P3 in magneto- and electroencephalography. Observers detected rare visual targets of a deviant shape and ignored rare non-targets of a deviant color. Source analysis was based on noise-normalized minimum-norm estimates in an individual, MRI-based source space. Critically, the cortical source space was extended to include the cornu ammonis of the hippocampus. Source analysis at the P3 peak showed strong sources in retrosplenial cortex and a weaker source in hippocampus. Activity in left somatosensory cortex was related to the button presses. Subsequent activity was observed in insula and anterior mid-cingulate cortex. Further simulation studies explored the spread of these sources and the plausibility of their combination to explain the data, and demonstrate that the combination of magneto- and electroencephalography increased the specificity of the source estimate. Overall, these data support an interpretation of the P3 as physiological indicator of activity in the limbic system for target and distractor stimuli.

## Introduction

The P3 or P300 (Sutton et al. 1965) is evoked in a multitude of contexts and has been one of the most studied electrophysiological brain responses in cognitive neuroscience (Polich 2020; Verleger 2020). Seeking the neural sources of the P3 with direct intracranial recordings from human epilepsy patients, P3-like activity was first found in the hippocampus (Halgren et al. 1980). Because resection of the hippocampus in such patients did not reduce the scalp P3, however, it was later concluded that the hippocampal P3 was generated in a closed source that does not significantly contribute to the P3 recorded with scalp electroencephalography (EEG) and with a maximum at electrode location Pz (Johnson 1988; Onofrj et al. 1992). Magnetoencephalography (MEG) studies that used dipole source analysis again reported that a main generator of the P3 was in the medial temporal lobe, supposedly in the hippocampus (Rogers et al. 1993; Tarkka et al. 1995; Basile et al. 1997). The focus of these studies was on the so-called P3b. This P3 variant is typically evoked by task-relevant stimuli, e.g. targets in a detection task (Hillyard et al. 1971), with a centro-parietal maximum at electrode location Pz. Another variant, P3a, is typically evoked by rare non-target or novel stimuli (Courchesne et al. 1975; Squires et al. 1975), often with a more frontal maximum, between electrode locations Cz and Fz. Reduced P3a was demonstrated in patients with unilateral medial temporal lobe lesions after posterior cerebral artery infarction, whereas P3b at electrode location Pz again remained unaffected in these patients (Knight 1996). Others proposed instead that several regions in the parietal and frontal cortex were more likely sources of both P3 variants in scalp EEG based on fMRI constrained source analysis (Bledowski et al. 2004; Mulert et al. 2004), intracranial recordings (Halgren, Baudena, Clarke, Heit, Liégeois, et al. 1995; Halgren, Baudena, Clarke, Heit, Marinkovic, et al. 1995), and lesion studies (Yamaguchi and Knight 1991), and the latter interpretation was more widely accepted until today, whereas the role of hippocampus remained controversial.

By applying noise-normalized distributed source analysis to combined MEG and EEG (M/EEG) data obtained with an auditory oddball paradigm, Das et al (2024) recently suggested that the main generator of the P3(b) is the retrosplenial cortex (RSC). RSC activity was similarly observed in fMRI for the same participants and could well explain the maximum of the P3 response in electrode location Pz in further simulation studies. While this finding would well explain that the parietal midline P3 was not reduced by loss of the medial temporal lobe, it does not necessarily imply that the hippocampal P3 cannot be observed in other electrode or sensor locations outside of the head, as well. Moreover, the RSC is tightly connected with the hippocampus (Kobayashi and Amaral 2003; Kobayashi and Amaral 2007), raising interesting interpretations of the P3 at a functional level. Medial temporal lobe activity was also observed when mapping the P3 with distributed source estimation, but the authors could not rule out that this activity could be caused by mislocalized source-estimate spread from the insular or auditory cortex (Das et al. 2024). The point-spread of such source models mainly depends on the proximity and alignment (source orientation) of dipolar sources in nearby brain regions, a well-known problem when interpreting the analysis of bioelectrical signals in MEG and EEG (Hämäläinen et al. 1993; Hauk et al. 2022). Leakage from other regions is important to consider in light of the fact that another generator of the P3 has been suggested in the insular cortex (Bledowski et al. 2004; Citherlet et al. 2020; Das et al. 2024) and because activity in the auditory cortex in the P3 latency range is readily evoked by auditory oddballs even without a task (Kretzschmar and Gutschalk 2010).

The present study sought to reexplore the limbic generators of the P3 with combined M/EEG and noise-normalized, anatomically constrained source analysis. Importantly, the individual cortical source space of the standard analysis stream was extended to cover the full hippocampus, including the cornu ammonis (CA), based on individually segmented MRI anatomy. This setup was used for the source analysis of experimental data and for simulation studies to probe their plausibility. The main hypothesis to be tested was that the CA regions of the hippocampus are a source of the P3 and can be recorded with M/EEG together with the RSC. In accordance with a previous study (Das et al. 2024), we expected that S1, insula, and anterior midcingulate cortex (aMCC) were further sources that contribute to the P3 complex. To this end, we used a visual instead of an auditory oddball paradigm, first, because it anatomically mitigates spread from auditory cortex to potential sources in the hippocampus and insula. For the same reason, white noise was used to mask the sounds of button presses in response to detected targets so that they would not activate auditory cortex. Second, visual stimuli were used to test if the source in RSC – found in previous auditory studies (Das et al. 2024; Doll et al. 2024) – was independent of the stimulus modality. The visual oddball paradigm included rare targets and distractors to explore if the source configuration can be transferred to rare stimuli that are not task-relevant. Finally, a number of simulations were added to this analysis, to probe the plausibility of these results, and provide an estimate for the spread and reliability associated with this source analysis.

## Materials and Methods

### Participants

Previous studies (Das et al. 2024; Doll et al. 2024) of the P3 with a similar recording setup had shown reliable group-level source activity in RSC with 12 and 14 data sets, respectively. To achieve at least a similar power for the source analysis, a total of 22 individuals (11 female, 11 male) participated in this study. None reported a history of neurological or ophthalmological disorders other than myopia. 20 participants were right-handed and two left-handed. All participants had either normal or corrected-to-normal vision, and none were color blind. Participants were students or employees recruited on campus, with no current neurological or psychiatric health issues or medication. Data from three participants were excluded due to measurement artifacts (n = 2) or co-registration errors (n = 1), such that the data of 19 participants were considered in the analysis presented below. The mean age of these 19 participants was 31 years (± 8 years standard deviation) and ranged from 24 to 61 years. Each participant gave written informed consent prior to participation, and the study was approved by the ethical council of the medical faculty at Heidelberg University (S-327/2016).

### Stimuli and experimental design

A visual oddball paradigm (Fig. 1) was utilized to evoke a P3 response. The paradigm consists of two similar experimental task conditions, which were used with the intention to reduce habituation in the relatively long recording session, and which were presented in the same order to all participants. In task 1, green circles (visual angle: 1.31° x 1.31°; radius = 3.5 cm), green heptagons (visual angle: 1.31° x 1.31°; 7 sides, side length = 3.37 cm, and apothem = 3.5 cm), and red circles (visual angle: 1.31° x 1.31°; radius = 3.5 cm) were presented as standard, target, and distractor stimuli. In Task 2, the standard stimuli were red squares (visual angle: 1.31° x 1.31°; radius = 3.5 cm), the target stimuli were red circles (visual angle: 1.31° x 1.31°; radius = 3.5 cm), and the distractors were green squares (visual angle: 1.31° x 1.31°; radius = 3.5 cm). All stimuli were presented with the same luminance and a refresh rate of 120 Hz in a dimly lighted environment (6.5 Lux). For each task, a total of 668 trials were presented, with the probabilities of standard, target, and distractor being 0.79, 0.12, and 0.09, respectively. Each stimulus lasted for 75 milliseconds and was presented at a regular interval of 2 s, with a jitter of ± 200 ms.

**Fig. 1.**
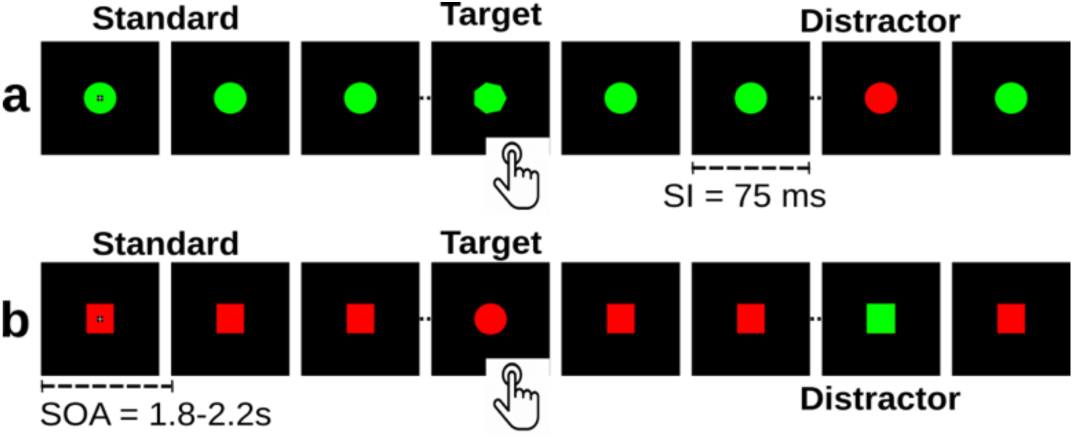
Overview of the experimental design. (a) Stimulus sequence in variant 1: green circles were presented as the standard stimuli, red circles as distractor stimuli, and green heptagons served as targets. (b) Stimulus sequence in variant 2: red squares were presented as standards, red circles as targets, and the green squares were distractors. Stimuli in both variants were shown in the middle of the screen against a black background. Each stimulus lasted for a stimulus interval (SI) of 75 ms and was presented with a stimulus-onset asynchrony (SOA) of 1.8 - 2.2 s. As soon as a target stimulus appeared on the screen, participants were asked to identify it by using their right index finger to click a response button.

All stimuli were created using Psychtoolbox-3 and presented on a black background with a resolution of 1920*1080 pixels. Images were back-projected on a transparent screen mounted in a window of a magnetically shielded room (height = 31 cm and width = 35 cm), 152 cm in front of the participants, with a BenQ TK700STi projector placed outside of the magnetically shielded room. The fixation cross was initially positioned precisely at the center of the screen. Participants were instructed to maintain their gaze on the fixation cross for the duration of each run. The participants were not instructed about the presence of distractor stimuli in the task. They were specifically told to identify only the target stimuli by clicking a response button using their right index finger as soon as a target stimulus appeared on the screen. First all trials of task 1 were presented, thereafter all trials from task 2; a short break was provided in between the two tasks. Prior to each task condition, a practice session consisting of 20 trials without any distractors was undertaken to allow participants to become familiar with the task. During the experiment, constant Gaussian white noise was presented at 80 dB SPL to mask the sound produced by pressing the response button. The sound was delivered binaurally using ER-3 earphones (Etymotics Research, Elk Grove Village, IL, USA) with foam tips.

### Data acquisition

The MEG recordings were conducted in a room equipped with four layers of magnetic shielding (IMEDCO AG, Hägendorf, Switzerland) during regular working hours. The recordings were performed using a Neuromag-122 whole-head system (MEGIN OY, Helsinki, Finland) comprising 61 dual-channel planar first-order gradiometers. The EEG data were obtained using a 64-channel equidistant cap (Easycap, Herrsching, Germany) where the electrodes extend far down from the low cheeks until below the inion. Cz was used as a reference electrode, and the impedances of all electrodes were kept below 3 kΩ. A bipolar electrode (near the clavicle) was utilized to record the ECG (electrocardiogram) and was later used as a marker to remove cardiac artifacts from the brain signals. EEG signals were amplified using two 32-channel Neuroscan amplifiers (Abbotsford, Victoria, Australia) and sampled together with the MEG at a frequency of 1000 Hz. The positions of MEG head coils, EEG electrodes, and some additional head-shape points (∼80 in total) were digitized together using the Polhemus Isotrack II digitizer (Colchester, Vermont, USA). Four head-position indicator coils were additionally digitized and used to acquire the participant’s head position relative to the MEG sensors.

Anatomical 3D-T1 (MPRAGE) MRI scans were obtained using a 3-T Siemens Magnetom scanner (Siemens Medical Systems, Erlangen, Germany) to create an accurate boundary-element head model. MPRAGE parameters included a field of view (FOV) of 256×256×192 voxels (voxel size = 1 mm^3^), repetition time (TR) = 1570 ms, echo time (TE) = 2.67 ms, inversion time (TI) = 900 ms, and flip angle = 9°.

### Data processing

Preprocessing and source estimation of MEG and EEG signals were carried out using MNE-Python (version 1.9; Gramfort, 2013). The raw data for each participant was visually inspected. Flat and noisy EEG channels (two to five) were interpolated using the spherical spline method (Perrin et al. 1989). The oversampled temporal projection (OTP) technique was used to remove flux jumps from MEG channels (Larson and Taulu 2018). The data were then filtered using a zero-phase finite-impulse-response (FIR) bandpass filter (0.5–30 Hz), and the EEG data were re-referenced to average reference. Next, the FastICA algorithm was used to remove cardiac artifacts and eye blinks (Hyvarinen 1999). Epochs were obtained using time intervals of 100 ms pre-stimulus to 1 s post-stimulus. Three stimulus-driven epoch conditions were constructed during this stage: standard, target, and distractor. A separate time window of 500 ms pre-stimulus and 500 ms post-stimulus was considered to create epochs to track the response-locked brain activity. Epochs involving missed trials and false alarms were excluded from the analysis. An automatic data-driven ‘autoreject’ method (Jas et al. 2017) was used to repair or exclude artifact-contaminated epochs. Prior to performing source analysis, the number of epochs across experimental conditions was equalized and averaged to produce separate evoked signals per condition using the ‘mintime’ scheme in MNE-Python.

### Cortex and Hippocampus surface reconstruction

Brain volume segmentation and cortical surface reconstruction for each participant were performed based on the high-resolution T1-weighted structural MRI scans using the standard stream of FreeSurfer (version 7.4.1), and individual cortex reconstructions were morphed to the spherical space of the fsaverage brain (Dale et al., 1999; Fischl, 2012). Additionally, individual participants hippocampal surfaces were derived by segmenting the bilateral hippocampus volume from each subject’s T1-weighted MRI with FreeSurfer (Fischl, 2012). Hippocampal surfaces include subregions CA1, CA2/CA3, and CA4 on the dorsal side; the subiculum and presubiculum on the ventral side; and a little portion of the hippocampal tail, including gray and white matter on both sides (Fig. 2). Topological defects such as surface-mesh distortion and vertex overlap were corrected, and surface smoothing was performed to achieve anatomically accurate surfaces suitable for M/EEG source modeling. The outer boundary of the segmented gray matter was treated as the hippocampal white-gray-matter boundary, equivalent to the white surface used in the cortex segmentation, enabling consistent surface-constrained dipole modeling in both compartments. Dipole orientations were constrained to be orthogonal to the local hippocampus surface, considering the laminar alignment of pyramidal cell populations (Murakami and Okada, 2015; Palomero-Gallagher et al., 2020) as the dominant generators of the hippocampal field potentials. For group-level analyses, individual hippocampal surfaces were transformed into spherical surface representations and aligned with the fsaverage hippocampus template via a non-linear spherical registration method. Based on the hippocampal subfields, overlay matching, and orthogonal dipole orientations relative to the local surface (dorsal or ventral side), two anatomical hippocampal surface labels were created. The first label corresponds to the dorsal hippocampal surface, which includes CA1–CA3 and the upper part of the hippocampal tail. The second label covers the ventral subiculum, presubiculum, and the ventral segment of the hippocampal tail.

**Fig. 2.**
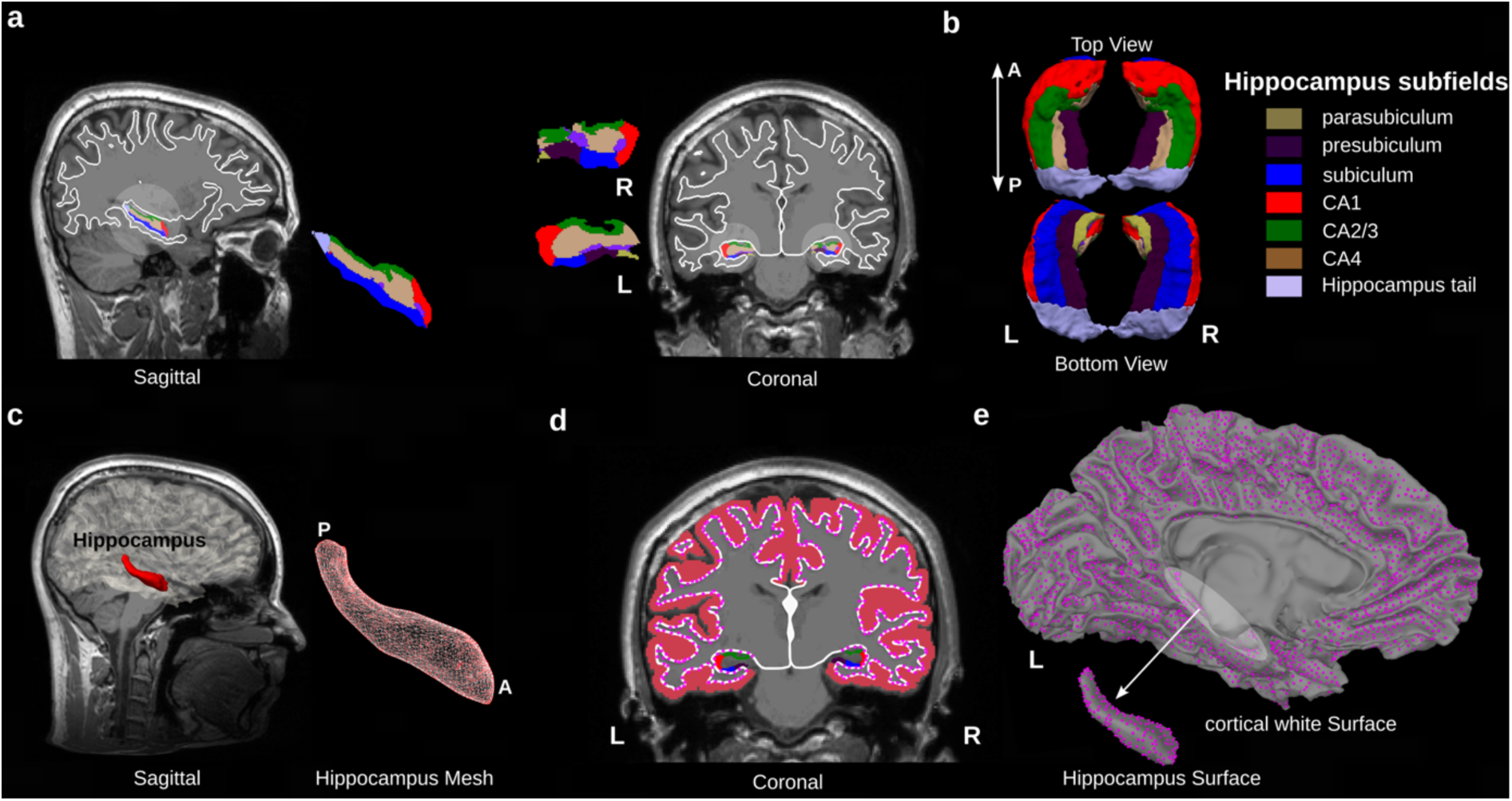
Hippocampus surface reconstruction and source space. (a) The hippocampus is segmented like a nucleus at the border between the surrounding white matter and the inner gray matter (colored subfields). In contrast, cortex is segmented at the outer border of the subcortical white matter (white line). (b) In an exemplary 3D reconstruction of the complete hippocampus, the different subfields are segmented and visualized from top and bottom; this view is referenced as dorsal and ventral hippocampus in the text, respectively (Iglesias et al. 2015). (c) The 3D hippocampus surface is aligned to the cortex surface. (d) In the combined cortical and hippocampal source space, the subiculum and parasubiculum are doubled and are therefore masked in the cortical source space, together with all midline dipoles that are unrelated to cortical structures. The remaining dipoles are indicated as magenta-colored dots. In the 3D hippocampus maps, dipoles in regions CA1-3 are mostly comprised in the dorsal hippocampus view, whereas subiculum and parasubiculum dipoles are comprised in the ventral hippocampus view, the latter of which is not shown in subsequent figures. Dipoles are orthogonal to the surface and per definition point outwards from the cortex segmentation and inwards to the hippocampus segmentation. (e) Combining the two sub source spaces allows for the analysis within a continuous, unified cortical-hippocampal source space. Directions are indicated as L (left), R (right), A (anterior), and P (posterior).

### Source localization

The cortical white surface and the outer hippocampal surface were aligned in three-dimensional space and integrated such that the vertices and faces of the hippocampal surface extended those of the white surface in the medial temporal lobe. This extended white surface was then utilized to establish a unified source space for cortical and hippocampal sources. Participant-specific source spaces were created by placing at least 10,000 sources (dipoles) per hemisphere at the gray-white matter boundary, with an average spacing of 3 mm (approx.) between the neighboring sources. Sources within the corpus callosum, and lateral ventricle wall were omitted (masked) during forward estimation using Human Connectome Project (HCP) parcellation (Glasser et al. 2016), as these regions do not include any part of the cortex. Similarly, dipoles in the subiculum complex of the cortical sub-source space were masked, because it overlapped with the ventral hippocampus sub-source space, and would then have been doubled. The dipole orientation was generally from the white-gray matter boundary towards the cortical surface, i.e. outwards for the cortex sub-space and inwards for the hippocampus sub-space. For inverse modeling, three-compartment boundary-element models (BEM) were used to create participant-specific, realistic head models (Hämäläinen and Sarvas 1989). The conductivities of the brain, skull, and skin were assumed to be 0.3 S/m, 0.006 S/m, and 0.3 S/m, respectively. Sources were assigned a loose orientation constraint (parameter value of 0.2), indicating a nearly perpendicular orientation to the cortical surface, which has been shown to improve source localization (Lin et al. 2006). A noise-covariance matrix was computed using the shrinkage technique from the 100-ms long pre-stimulus interval (Engemann and Gramfort 2015). Combined M/EEG source estimation was performed using the linear L2 minimum-norm estimator (MNE) (Hämäläinen and Ilmoniemi 1994; Molins et al. 2008) with noise normalization (Dale et al. 2000), also known as dynamic statistical parametric mapping or dSPM. Depths weighting (0.8) and regularization (λ = 0.111) parameters were set to the suggested default values (Lin et al. 2006; Gramfort et al. 2013). For the group-level analysis, all individual dSPMs were transformed into the fsaverage space. For tasks 1 and 2, separate dSPMs were computed to account for different head positions during MEG recordings. Since these separate dSPMs produced very similar results, an average across tasks was performed to create a single dSPM solution. Later, this solution was used to show significant brain activity maps (p < 0.01; fixed effects).

To extract local source waveforms, several regions of interest (ROI) were defined. The ROIs were defined as such that the polarity within regions was expected to be uniform, i.e. they were limited to one bank of a sulcus, where the actual anatomical source was expected (cf. Das et al., 2024 for details). In total, 5 bilateral ROIs were used, including the left and right insula, somatosensory hand area (S1), cornu ammonis (CA), retrosplenial cortex (RSC), and anterior midcingulate cortex (aMCC). Insula and RSC were directly adopted from the previous study (Das et al. 2024). The CA ROI was not part of the previous analysis and was based on the dorsal surface of the hippocampus (cf. Fig. 2). The S1 and aMCC ROIs were adjusted slightly in comparison to Das et al., 2024 in order to more accurately include the activation peak of the current data set. Why the activation peaks of S1 and aMCC did not match between studies remains unclear; one possible explanation for S1 could be the utilization of different response devices. For each experimental condition, dSPM-based source time-courses for these ROIs were calculated separately by averaging the activity across all sources within each ROI using the ‘mean-flip’ method implemented in MNE-Python (Gramfort et al. 2013). This method compensates for signal cancellation by correcting the polarities of the dipoles in comparison to the average orientation produced by all dipoles for a given ROI. This process was carried out for each participant separately before producing a grand-average source time-course across the participants. Because of the restriction of our ROIs to one bank of a sulcus or gyrus, the application of the mean-flip procedure produced only minor effects and did not alter our conclusions.

Peak-latency analysis was performed on the ROI-based source time-courses. For this procedure, the evoked signals were band-pass filtered at 1–8 Hz using a zero-phase-shift FIR filter. Peak latencies for individual ROIs were estimated within a time window between 300-700 ms after stimulus onset. Afterwards, two-sample t-tests were carried out to compare peak latencies between ROIs for the target and distractor P3. The p-values of the tests were corrected for multiple comparisons using a Bonferroni correction with an alpha level of 0.05, and only the corrected p-values (p<0.05) are reported.

### Cortical source simulations

A simulation model was implemented to probe how well the experimental data could be explained by a sparce source model (2.8.), and to estimate sensitivity profiles and the spread between brain regions (2.9). For each participant, bilateral anatomically constrained sources were simulated based on the selected ROIs. All dipolar sources within each ROI were activated using a half-sinusoid with a base frequency of 10 Hz. The scaling of the source currents for dipoles was adjusted to ensure that the absolute mean value of all dipoles within an ROI equaled 25 nAm at the peak activity, corresponding to the time instant with the absolute maximum amplitude. This ensures uniform source strength across ROIs, regardless of variations in their sizes. The polarity of the simulated source time-course was also adjusted with the observed polarity in the source analysis. Finally, dipoles within the left and right hemisphere ROIs were combined to create a bilateral source. This process yielded four bilateral sources: insula, CA, RSC, and aMCC, with the exception of the S1 source, where only left-hemisphere dipoles were considered to replicate contralateral, response-hand-specific brain activity. The simulated sources were then transformed back to the electrode/sensor space to produce simulated evoked signals by multiplying the simulated time-series of the sources by the gain matrix. Sensor-level noise was simulated by scaling multivariate gaussian white noise based on individual noise-covariance matrices estimated from the original M/EEG recordings of each participant, assuming 200 averaged trials.

### P3 pattern analysis using simulated sources

The activation patterns of anatomically constrained simulated sources were linearly combined to explain the P3 peak activation in experimental M/EEG data for targets and distractors. The aim of this procedure is to find a simulated activation pattern such that, when the simulated sources are multiplied by their respective weights and linearly combined, they explain the actual P3 pattern in a least-squares sense. A multiple linear regression approach (Haufe et al. 2014) was utilized here to optimize the weights from the simulated sources. This model can be expressed as:

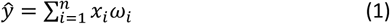

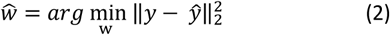

Where, ŷ is the predicted simulated P3 pattern; *y* is the actual P3 pattern; *x_i_* is the activation pattern of the *i*^th^ simulated sources; ω_i_ is the weight of the *i*^th^ simulated source; and *ŵ* are the optimal weights.

First, evoked M/EEG data were band-pass filtered at 1–8 Hz using a zero-phase FIR filter, and a search window of 300–700 ms was used for reliable P3 peak detection. MEG data were transformed into 61 virtual magnetometers (Gramfort et al. 2013) to provide a similar number of MEG sensors and EEG electrodes, and a more similar sensor-level signal-to-noise ratio. P3 activation peaks were then extracted from Pz across participants to compute a grand average P3 pattern. Later, the grand-average P3 distribution is used as the target pattern (dependent variable) in the regression model. Five simulated sources (insula, S1, CA, aMCC, and RSC) were considered as the input variables (input features). Since EEG and MEG data consist of different units, z-scored standardization was performed across individual sensor types utilizing a 50-ms pre-stimulus time interval. We performed this step for both input and actual target variables, ensuring that we could compute latent weights by combining EEG and MEG data, as long as the data remained z-scored (unit-less). A multi-collinearity test was also performed to establish linear independence across input variables (simulated sources). A variance-inflation factor of less than 5 was observed across simulated sources during the test, meaning all the simulated sources were fairly independent from each other. Finally, to quantify the performance of the model, residual variance was calculated from the *R*^2^ score of the model, which was calculated between the actual and predicted P3 pattern. This step was separately performed for EEG and MEG data. The optimized latent weights were multiplied by the source strength (25 nAm) of the simulated sources to provide a rough estimate of the source strength, leading to the generation of the actual P3 peak pattern in MEG and EEG.

### Source leakage and source sensitivity

Spatial source leakage is inherent for MEG and EEG source signals due to their limited spatial resolution and the non-uniqueness of the inverse problem (Sarvas 1987). The degree to which source leakage varies depends on the type of source estimation method being used, the sensor configurations used, source depth, and source orientation (Hauk et al. 2022). In this study, potential leakage from and into the selected ROIs with the dSPM were characterized using the concept of point-spread functions (PSFs) to identify source leakage. The dSPM-based resolution matrix (Samuelsson et al. 2021) was computed for each participant based on simulated sources. PSFs for each ROI were extracted and normalized by its maximum peak activity. Then, an average group of ROI-specific PSFs across participants was performed. Finally, an absolute Pearson correlation test was performed between the leakage activation (grand-PSFs) of one ROI and the rest of the ROIs to determine how the point-source activity from one ROI would spread to other ROIs. Correlation values range from 0 (no leakage) to 1 (complete spatial overlap). Conversely, to quantify the sensitivity of the recoding methods for the ROIs used, the signal-to-noise ratio (SNR) within the ROIs was calculated based on the simulated M/EEG activity at the source level. For each subject and ROI, the maximum source amplitude (*a*) was extracted as peak activity from the reconstructed source time-course and used to compute the SNR in dB (Goldenholz et al. 2009):

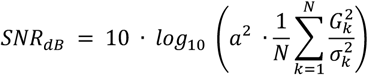

where *G_k_* is the forward model gain at sensor *k*, *σ_k_^2^* is the noise variance at that sensor, and *N* is the total number of sensors. SNR was computed separately for MEG, EEG, and combined M/EEG for each subject. Note that this estimate is based on weighted MNE (and not dSPM).

## Results

The data are based on a visual oddball paradigm with two non-standard stimuli. The targets were of the same color as the standards but of a different shape, and the distractors were of a different color. Participants were instructed to press a response button for the target stimuli, and to ignore the non-standard distractor stimuli. Two variants of the stimulus paradigm were used (Fig. 1) with the intention to reduce response habituation. The hit rates for the two variants were similar and not significantly different across participants, with hit rates of 99.8 ± 0.67% in variant 1 and 99.3 ± 0.71% in variant 2 (mean ± standard deviation). The reaction times were 423 ± 49 ms and 425 ± 60 ms (mean ± standard deviation) in variants 1 and 2, respectively. As the separate analyses of the M/EEG data were also highly similar, the two variants were combined for the analysis presented below, to yield a better signal-to-noise ratio. The separate results for task 1 and task 2 are shown in the appendix (Fig. S1).

In the grand average scalp-EEG waveforms (Fig. 3, b; left panel), four subsequent peaks are observed in response to the standard stimuli, with response latencies around 100, 200, 300, and 390 ms. The target response (Fig. 3a) was instead dominated by a strong P3 with a response latency of 396 ± 50 ms (mean ± standard deviation, measured at electrode Pz). The third and fourth peaks were also enhanced for the distractor stimulus (Fig. 3b), but much less than for the target stimuli. We consider the fourth peak with a mean latency of 383 ± 80 ms (measured at electrode Pz) as the distractor P3 in this case. Similar responses were observed in the MEG sensor-level waveforms (Fig. 3; right panel), but with somewhat less distinct P3 compared to EEG. The waveforms of the evoked response averaged relative to the time of the button responses (Fig. 3c) showed peaks shortly after the button press, which were more broad-based in EEG (left) compared to MEG (right).

**Figure 3.**
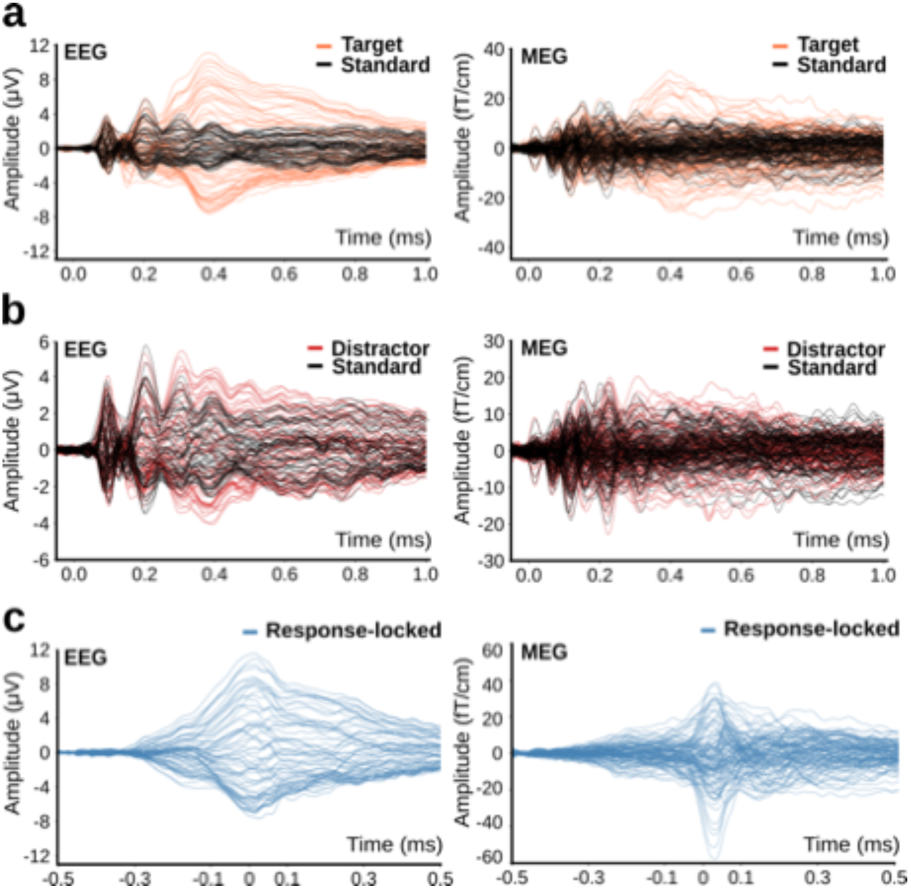
Grand-average evoked signals for EEG and MEG (N=19). (a) Stimulus-locked evoked signals for EEG (left) and MEG (right) for targets (coral) and standards (black). Only the targets evoke prominent P3 activity, which occurs mostly in the 350–450 ms time range. (b) The evoked responses for distractors (red) and standards (black) are shown for EEG (left) and MEG (right). There is only a relatively weak distractor P3 observed in the range of 300 – 500 ms. Note the different amplitude scaling in a and b. (c) Evoked signals for detected targets, averaged by the button press. The somatosensory feedback is evident in MEG (right), which shows a very pronounced and rapidly growing response after the button press. In contrast, EEG (left) is dominated by a progressively increasing signal prior to the initiation of the button press.

Grand-average dSPM source maps of the combined M/EEG were then calculated in a time window of 325–475 ms after stimulus onset, covering the P3 peak across participants. The dSPM maps for the target stimuli (Fig. 4a) first showed negative polarity in the RSC, accompanied by positive activity in two spots along the (dorsal) posterior cingulate cortex (PCC), i.e. on the side opposite to RSC in the cingulate gyrus. As has been shown before and is demonstrated further below, such pairs of positive and negative-polarity on opposite sides of a gyrus (or sulcus) are typically observed in dSPM maps with sources on one side of a gyrus. Prominent negative-going activity was also observed in the dorsal hippocampus, comprising most of CA. The ventral part of the hippocampus segmentation (not shown), comprising the subiculum, presubiculum, and parasubiculum revealed positive-going activity, the continuation of which is seen in some spots along the hippocampal sulcus in the cortex maps. In the convexity of the parahippocampal gyrus, extending anterior into the entorhinal cortex and lateral up to the fusiform gyrus, activity is again negative going. At the border between the medial temporal lobe and RSC clusters, activity extends into the calcarine fissure. Some midline activity was finally observed in the right anterior cingulate gyrus (ACC). On the cortical convexity, the largest cluster of activity mapped is located in the insular cortex in both hemispheres. Less prominent spots of activity are mapped to the central region and the superior temporal sulcus. Mapping the activity evoked by standards in the same time range and with the same cutoff reveals almost no activity. Accordingly, the difference between targets and standards (Fig. 4a) reveals similar activity patterns in RSC and hippocampus, as observed for the targets. Focal activity in S1 is also similarly observed in target and difference maps. Insular activity is not significant in the difference maps of the P3 peak, but is significant in a later time window (Fig. S2) along with stronger insular activity. Similar source analysis results were obtained with the LCMV (linearly constrained minimum variance) beamformer (Fig. S3a) or sLORETA (standardized low resolution brain electromagnetic tomography; Fig. S3b).

**Fig. 4.**
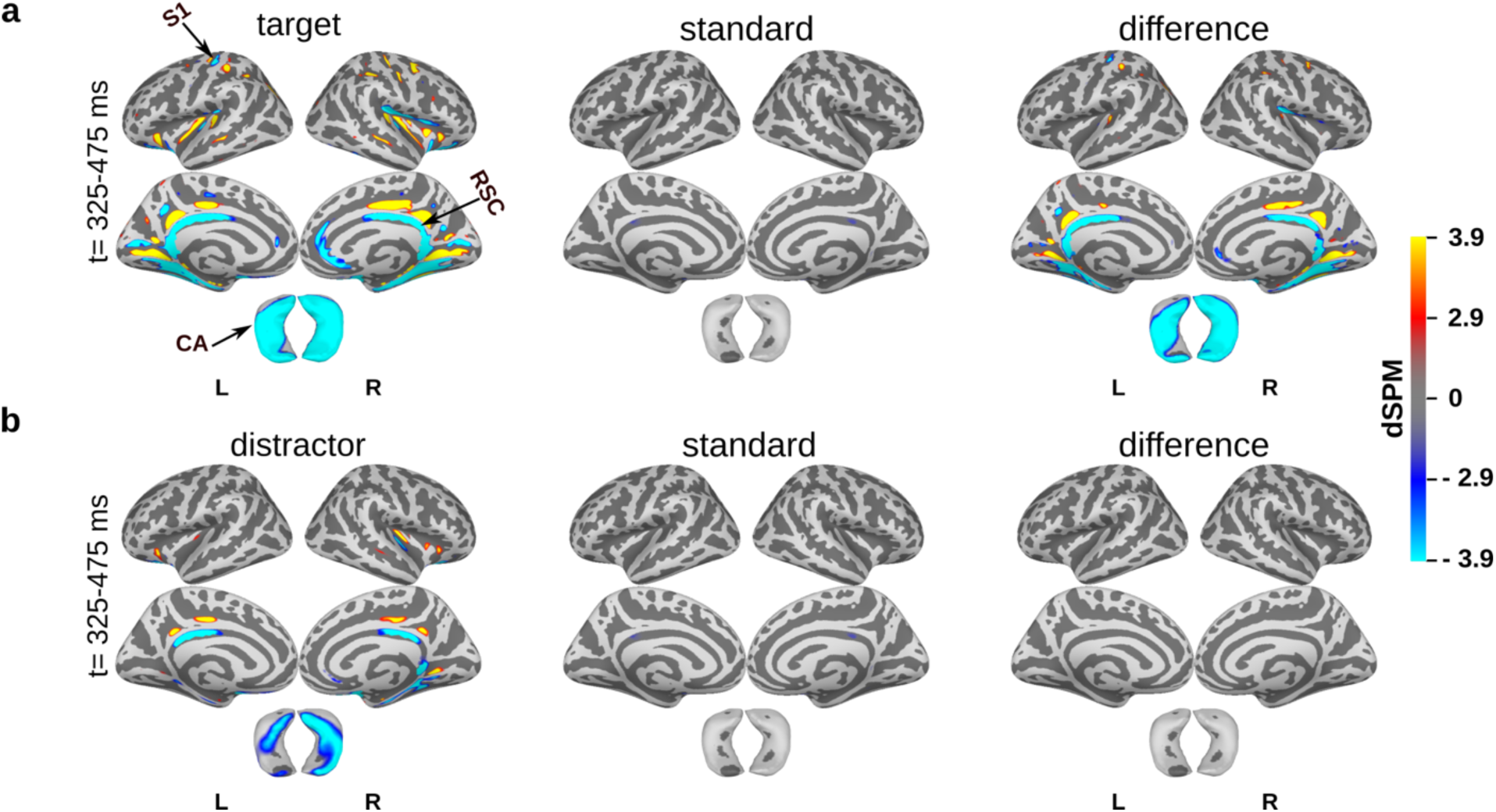
Grand-average dSPM activation maps for combined M/EEG (N=19). (a) Combined M/EEG dSPM maps (p<0.01) for targets (left), standards (middle), and the contrast between targets minus standards (right) in the time window 325–475 ms after stimulus onset. (b) dSPM maps for distractor stimuli, similarly arranged. While activity in RSC, insula, and CA is observed for distractors and not for standards, no significant difference is mapped between these two conditions (right column). The number of trials were equalized between targets, distractors, and standards prior to source estimation. Highlighted brain regions include retrosplenial cortex (RSC), primary somatosensory cortex (S1), and the cornu ammonis (CA). Note that only the dorsal view of hippocampus is shown and that for technical reasons the activity is plotted on the outside, while the polarity shown corresponds to the cortical surface in the inside of hippocampus (cf. Fig. 2).

For a comparison with a previously used auditory oddball paradigm (Das et al. 2024), a re-analysis including the hippocampal source space (Das 2025) has been added in Fig. S4: Activity in RSC extends somewhat more anterior and posterior along the cingulate gyrus in the present study (Fig. 4a), but completely includes the activity mapped in the auditory data set. Activity mapped in the insula also overlaps, but is more prominent in the auditory data, where it converges with activity in the auditory cortex, where no activity is mapped in the present, visual study. While CA activity mapped in the present study extends from the posterior to the anterior end of CA (Fig. 4a), it is only observed in the anterior half of CA for the auditory data (Fig. S4b). This is likely caused by an interaction with positive-going spread from the auditory cortex; since stronger activity is mapped in left auditory cortex, this would explain why stronger CA activity is mapped to the right, considering stronger cancelation in left CA. Activity in aMCC is better compared for the latency after 500 ms (visual: Fig. S2, auditory Fig. S4c), and extends somewhat more anterior in the visual data.

One intention for the present, visual study was to avoid the spread from auditory cortex to CA. To evaluate potential spread from occipital cortex and fusiform gyrus, visual source activity was mapped (FIg. S5). At the cutoff used to map the P3, no significant activity was mapped for the P100. In the subsequent time window (150 - 200 ms), focal activity was mapped to the border of lingual cortex and fusiform gyrus, but no significant spread to was observed CA at the same cutoff. It appears unlikely that later sensory activity from fusiform gyrus is stronger and spreads significantly to CA, but this cannot be certainly excluded since fusiform activity is also observed in the P3 time window.

The maps for visual distractor stimuli are shown in Fig. 4b: distractor maps show consistent activity in the bilateral RSC and hippocampus. Some spots of activity in the convexity are observed in insular cortex, also more prominently on the right. Neither of these activities are significant in the standard maps nor in the difference maps.

The M/EEG source time courses were further evaluated in ROIs covering RSC, CA, insula, aMCC, and S1 (Fig. 5): For targets (Fig. 5a), source waveforms revealed negative-going waves in bilateral RSC, CA, and aMCC. Only insula showed a positive-going wave, while a biphasic wave was observed in left S1. Peak latencies for targets (Fig. 6a) were earlier in RSC (443 ± 90 ms) and CA (427 ± 58 ms), compared to insula (574 ± 86 ms) and aMCC (598 ± 55 ms; mean ± standard deviation). The response to distractor stimuli (Fig. 5b) also showed long-latency peaks in the P3 range in RSC (391 ± 73 ms), insula (537 ± 79 ms), and CA (425± 68 ms). Compared to the target P3, however, the response to the distractors appears more similar to the standard stimuli in amplitude. ROI latencies are summarized in Fig. 6; P3 latencies in standard, midline electrode positions are shown in Fig. S6 for comparison. An analysis based on the data averaged to the button presses is summarized in Fig. S7.

**Figure. 5.**
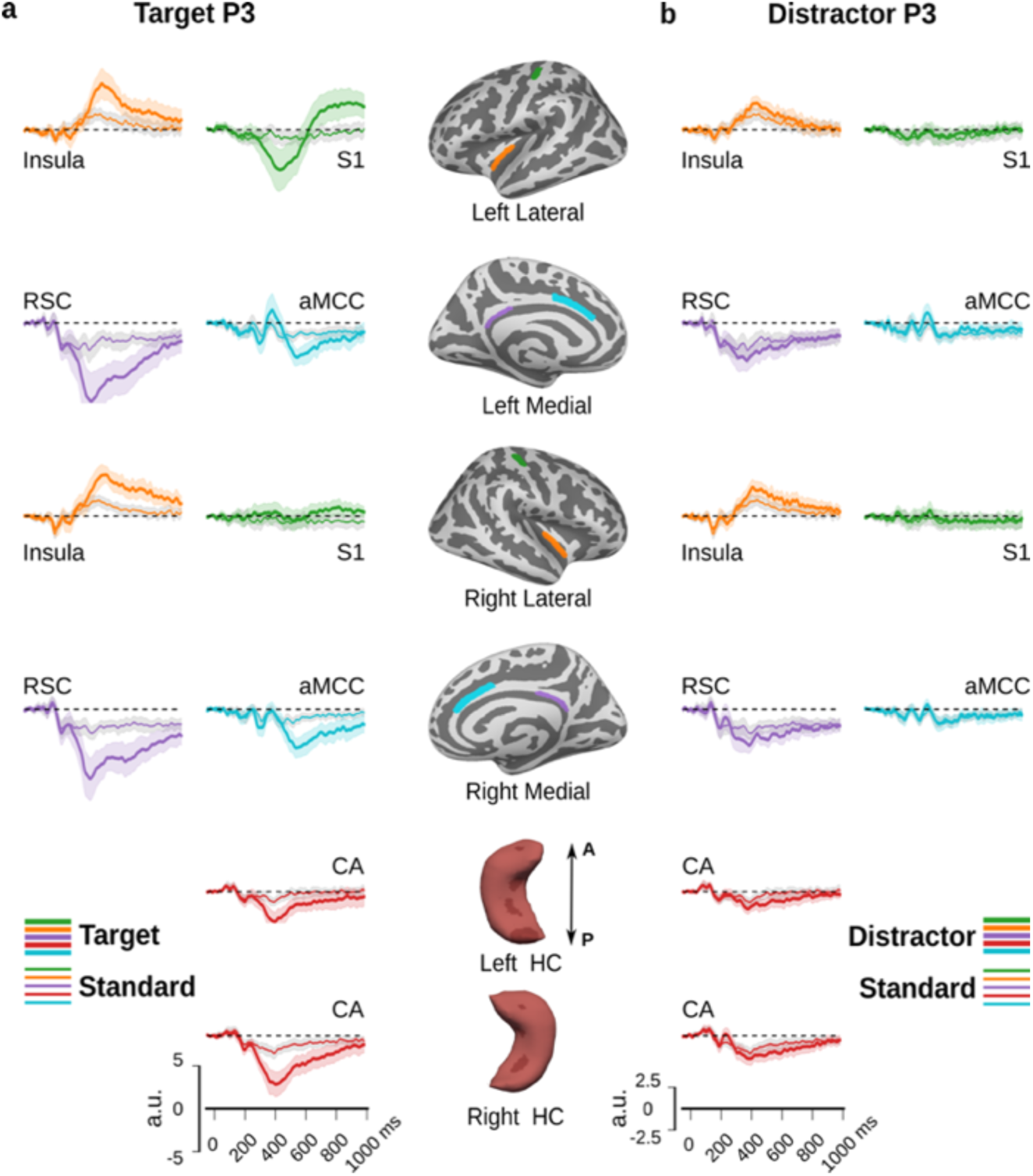
ROI-based M/EEG source waveforms for target, standard, and distractor (N=19). (a) M/EEG source waveforms extracted from dSPM estimates across participants for targets (thick lines) and standards (thin lines). The ROIs are plotted in the inflated fsaverage cortex and hippocampus (HC) in the middle column. The same color code is used to represent each ROI and its corresponding source waveform. The ROIs include the insula, primary somatosensory cortex (S1), retrosplenial cortex (RSC), anterior midcingulate cortex (aMCC), and cornu ammonis (CA). The shaded areas represent the 95% confidence intervals, calculated using a non-parametric bootstrap method. (b) Similar arrangement of source waveforms but for distractors (thick lines) and standards (thin lines).

**Fig. 6.**
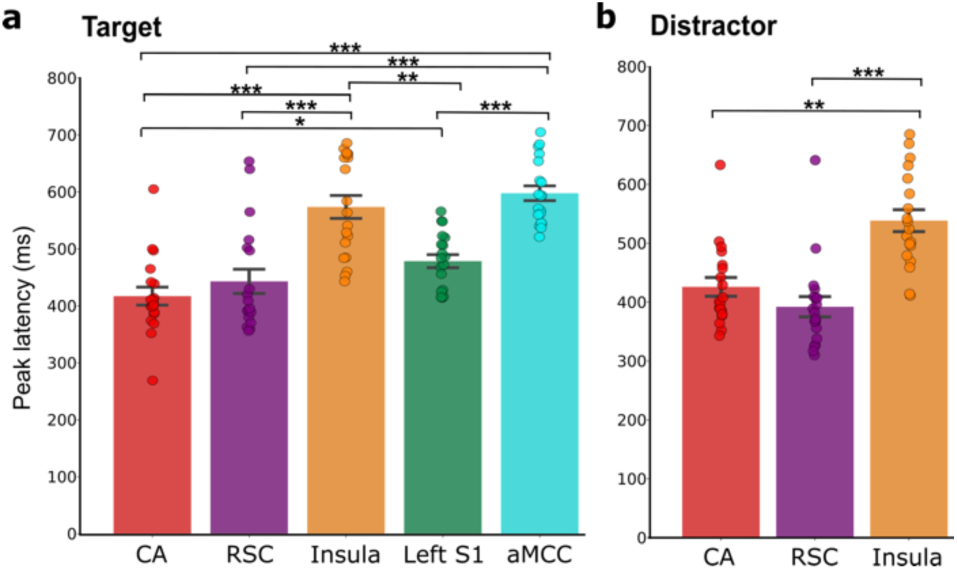
Peak-latency analysis of ROI-based source time courses (mean ± standard error; N=19) for (a) targets and (b) distractors P3. The ROIs include the cornu ammonis (CA), retrosplenial cortex (RSC), insula, primary somatosensory cortex (S1), and anterior midcingulate cortex (aMCC). Significant differences between peak latencies for ROIs are indicated by asterisks (* p < 0.05, ** p < 0.01, *** p < 0.001) based on t-tests for repeated measures with Bonferroni correction for multiple comparison.

Next, to estimate the plausibility and relative contribution of these sources modeled as ROIs, the MEG sensor and EEG scalp distribution were modeled with a linear combination of simulated activity based on the ROIs (Fig. 7a). To reduce the degrees of freedom, this sparse model was constrained to equal source amplitudes in the left and right hemispheres for each pair of ROIs, with the exception of S1, which was only modeled in the left hemisphere. When this model was applied to the peak of the target P3 (Table 1), the strongest weighting was applied to RSC, followed by CA with about a third of the signal strengths. Activity in S1 and aMCC was yet lower, and insula received no weighting at all when all sources were used. Leaving out single sources in this model revealed that RSC explained the most variance in EEG, whereas S1 explained the most variance in MEG. The second most variance in EEG was explained by S1 and in MEG by CA. Matching the overall lower residual variance in EEG compared to MEG, the maps resulting from this simulation are more similar to the original data in EEG than in MEG (Fig. 7b), although the general distribution can be modeled in both modalities. A similar pattern was observed when the model was applied to the distractor P3 (Fig. 7c). The absolute amplitude of the EEG and MEG in this simulation depends critically on the head model. Because the head model applied here does not include a separate cerebro-spinal fluid (CSF) compartment, the EEG amplitude is likely overestimated (Piastra et al., 2021). This can be compensated for by proportionally decreasing the conductivity of the skull, instead (Stenroos and Nummenmaa 2016). When the simulation is based on such a modified head model (Fig. S8), the estimated source strength is higher to explain the recorded EEG, in particular in the RSC and CA (Table S1). Note that the dSPM estimates used in the rest of this paper are robust to modifications of skull conductivity (Fig. S9), matching previous evaluations (Stenroos and Hauk 2013). Since the combination of MEG and EEG is based on the noise normalized data for the dSPM estimate (Liu et al. 2002), the exact physical scaling of MEG and EEG can be neglected here.

**Fig. 7.**
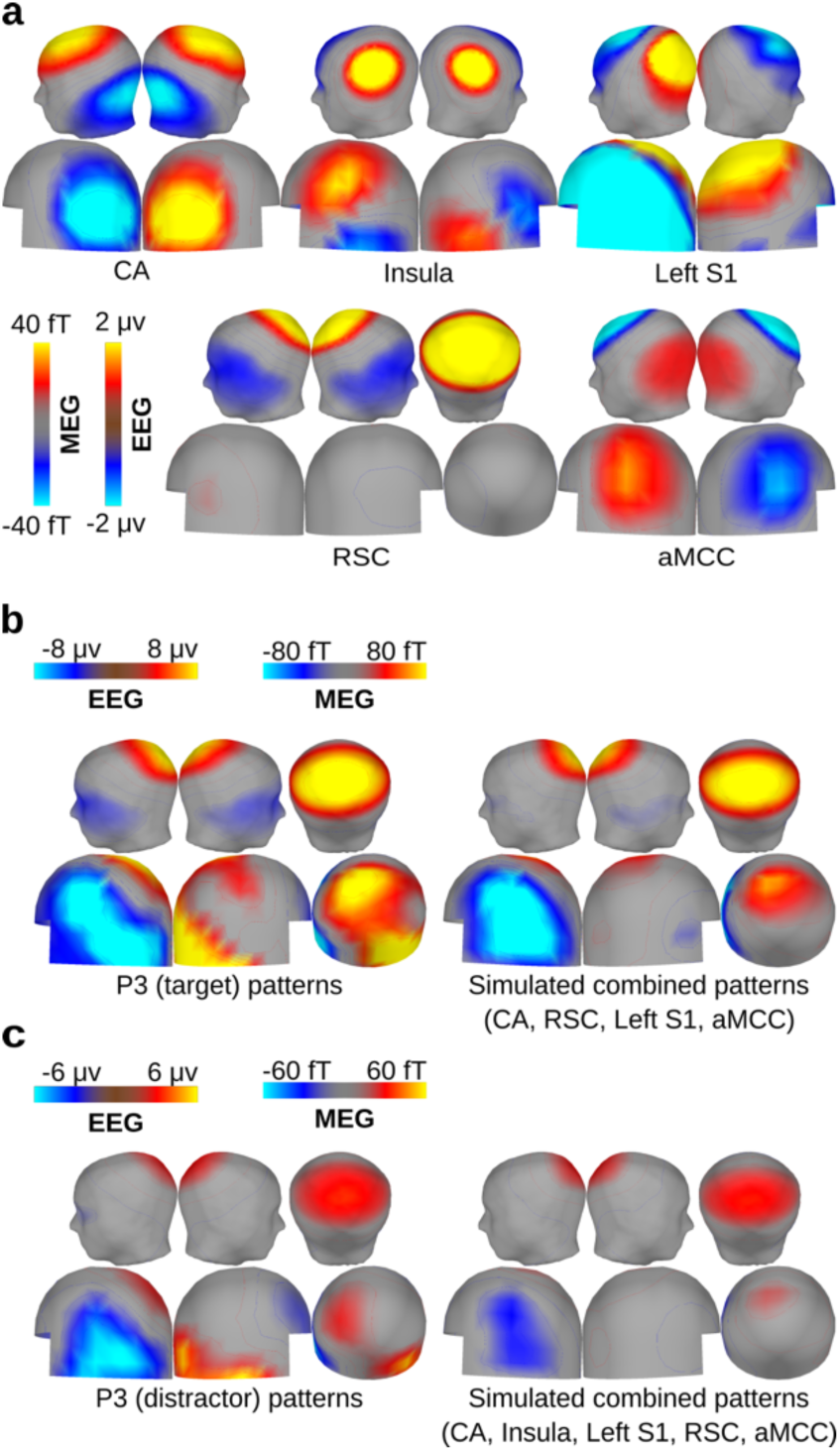
Simulated M/EEG sources and sparse P3 source model. (a) Spatial pattern of scalp EEG (top) and virtual magnetometer maps (bottom) for MEG, derived from simulated sources (grand average, N=19) in the bilateral cornu ammonis (CA), insula, left primary somatosensory cortex (S1), bilateral retrosplenial cortex (RSC), and anterior midcingulate cortex (aMCC). (b) Grand-average (N=19) scalp EEG and virtual MEG magnetometer maps at the peak latency for the P3 in Pz for targets (left). Weighted (cf. table 1) combination of simulated sources in CA, insula, S1, RSC, and aMCC that best explain the P3 for targets (right). (c) Same arrangement as in b for distractor-evoked P3.

**Table 1:** Relative weights and residual variance for the sparse model of the grand-average M/EEG for target P3 and distractor P3 with simulated, ROI-based sources. CA, cornu ammonis; RSC, retro-splenial cortex; S1, primary somatosensory cortex; aMCC, anterior middle cingulate cortex; EEG, electroencephalography; MEG, magnetoencephalography.

| Condition | Simulated source activation strenght (nAm) |  |  |  |  | Residual variance (%) |  |
| --- | --- | --- | --- | --- | --- | --- | --- |
|  | CA | RSC | Insula | left S1 | aMCC | EEG | MEG |
| Target P3 | 22.0 | 63.5 | 0.0 | 14.5 | 10.3 | 7.0 | 33.4 |
|  | 22.0 | 63.5 | - | 14.5 | 10.3 | 7.0 | 33.4 |
|  | 13.8 | 66.5 | 7.0 | 15.0 | - | 6.6 | 34.7 |
|  | - | 71.0 | 5.5 | 15.3 | 0.0 | 7.0 | 40.8 |
|  | 42.8 | - | 0.0 | 13.0 | 22.0 | 69.2 | 34.8 |
|  | 31.3 | 56.8 | 0.0 | - | 18.5 | 13.9 | 98.0 |
| Distractor P3 | 7.8 | 29.3 | 0.5 | 5.5 | 7.3 | 8.2 | 46.3 |
|  | 8.3 | 29.3 | - | 5.5 | 7.5 | 8.3 | 46.1 |
|  | 10.3 | 28.8 | 0.0 | - | 11.3 | 9.7 | 99.6 |
|  | - | 32.5 | 5.5 | 5.8 | 0.0 | 7.4 | 52.7 |
|  | 1.5 | 31.8 | 5.8 | 5.8 | - | 8.2 | 49.7 |
|  | 23.3 | - | 0.0 | 5.0 | 16.0 | 72.3 | 41.0 |

To further evaluate the individual contribution of MEG and EEG, single-modality source estimates of the target and distractor conditions were calculated (Fig. 8). This analysis demonstrates that - at the same cutoff as in Fig. 4 - RSC activity is not observed in MEG, but only in EEG, matching the contribution of the sparse source model (Table 1). Some activity in right CA is also observed in MEG, but again the activity is more prominent in EEG. In the EEG maps, the distribution of activity is also wider. This is partly an effect of stronger sources, and accordingly reduced by a higher cutoff, but it may additionally be caused by a sharpening of the inverse in the combined M/EEG source analysis.

**Fig. 8.**
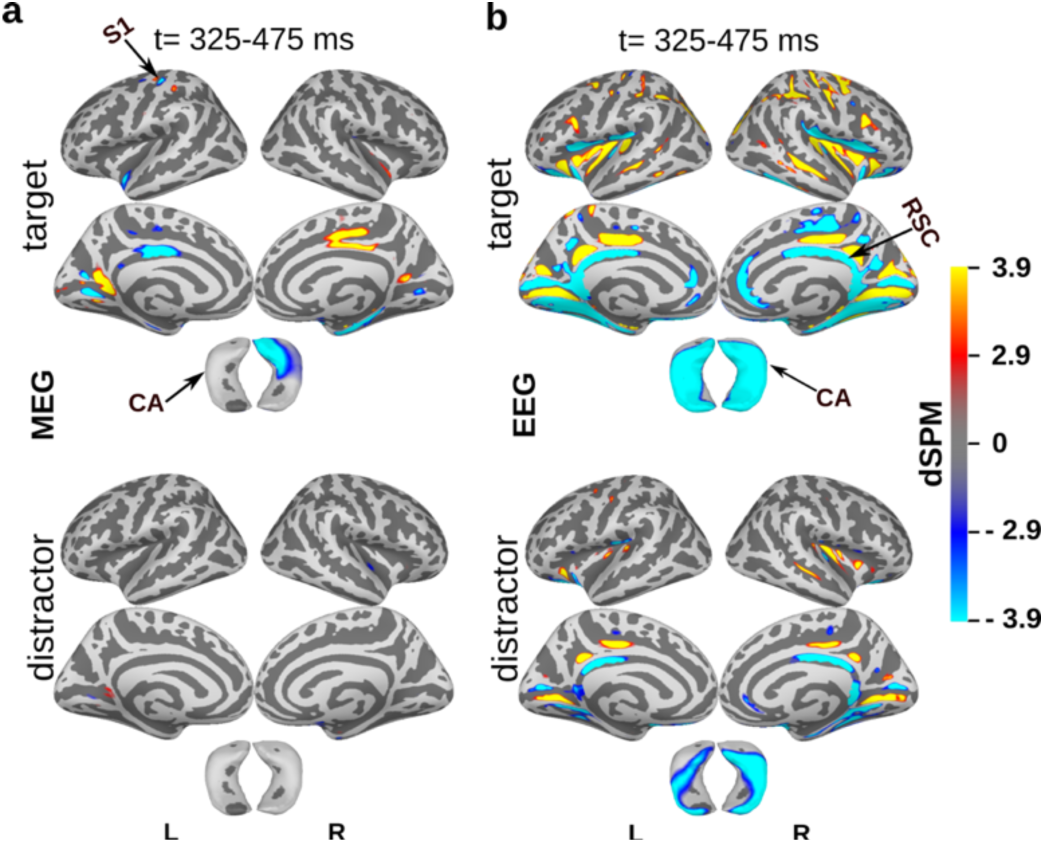
Grand-average dSPM activation maps calculated separately for MEG and EEG. (a) Separate MEG and (b) EEG dSPM maps (p<0.01; N=19) for visual target (upper) and distractor stimuli (lower) in the time window 325-475 ms after stimulus onset. Highlighted brain regions include retrosplenial cortex (RSC), primary somatosensory cortex (S1), and cornu ammonus (CA).

To support the interpretation of the dSPM maps and to better dissociate “real” activity from source spread, we computed simulated M/EEG activity based on single-region, bilateral ROIs (except left S1) and applied dSPM source analysis. (Note that the scaling is normalized for the activity within the ROI used for the simulation and therefore differs between MEG, EEG, and M/EEG maps, respectively). These maps reproduce several aspects of the dSPM source analysis shown in Fig. 4: In particular, activity in CA spreads to the para-hippocampal gyrus and to the calcarine fissure. Spread to the calcarine fissure is even more prominent when the simulation is made only for MEG or EEG; it is noteworthy that the spread observed for CA simulation shows different polarity patterns in the calcarine fissure for MEG and EEG, respectively, and that this pattern reproduces the pattern observed in the single-modality source analysis of the target condition (Fig. 8). The similarity of this pattern for visual (Figs. 8) and auditory targets (Fig. S4) in M/EEG further supports that activity mapped around the calcarine fissure represents spread from CA rather than local activity. Prominent spread from CA is also observed in the insula. Conversely, spread from the particular insular ROI used here is mostly observed in anterior and lateral, more radially oriented parts of CA. RSC spreads to the PCC, and a little to the calcarine fissure. In the EEG-only maps, some spread from RSC is also observed in the bilateral central sulcus and in the intraparietal sulcus, where activity is similarly observed in the EEG-only analysis (Fig. 8b). Finally, spread from S1 to the posterior MCC is observed in MEG, and may explain at least part of the strong left negative-, right positive-going activity observed in the MEG-only analysis (Fig. 8a).

Critically, these maps as well as the correlational analysis of spread between ROIs (Fig. 9b and Fig. S10-S12) demonstrate that the combined M/EEG maps are usually less affected by spread between ROIs compared to both, the MEG- and the EEG-only analysis. Among the 10 ROI interactions evaluated, there are two exceptions from this observation: The spread between insula and CA is stronger in MEG (0.53) and M/EEG (0.38) than EEG (0.24). This is of interest, because the spread between CA and insula is also the strongest of all combinations tested, probably caused by their spatial proximity. Second, the spread between aMCC and insula is stronger in M/EEG (0.12) and EEG (0.18) than MEG (0.05) alone. There is no condition in which spread in M/EEG was stronger than in both single modalities, however.

**Fig. 9.**
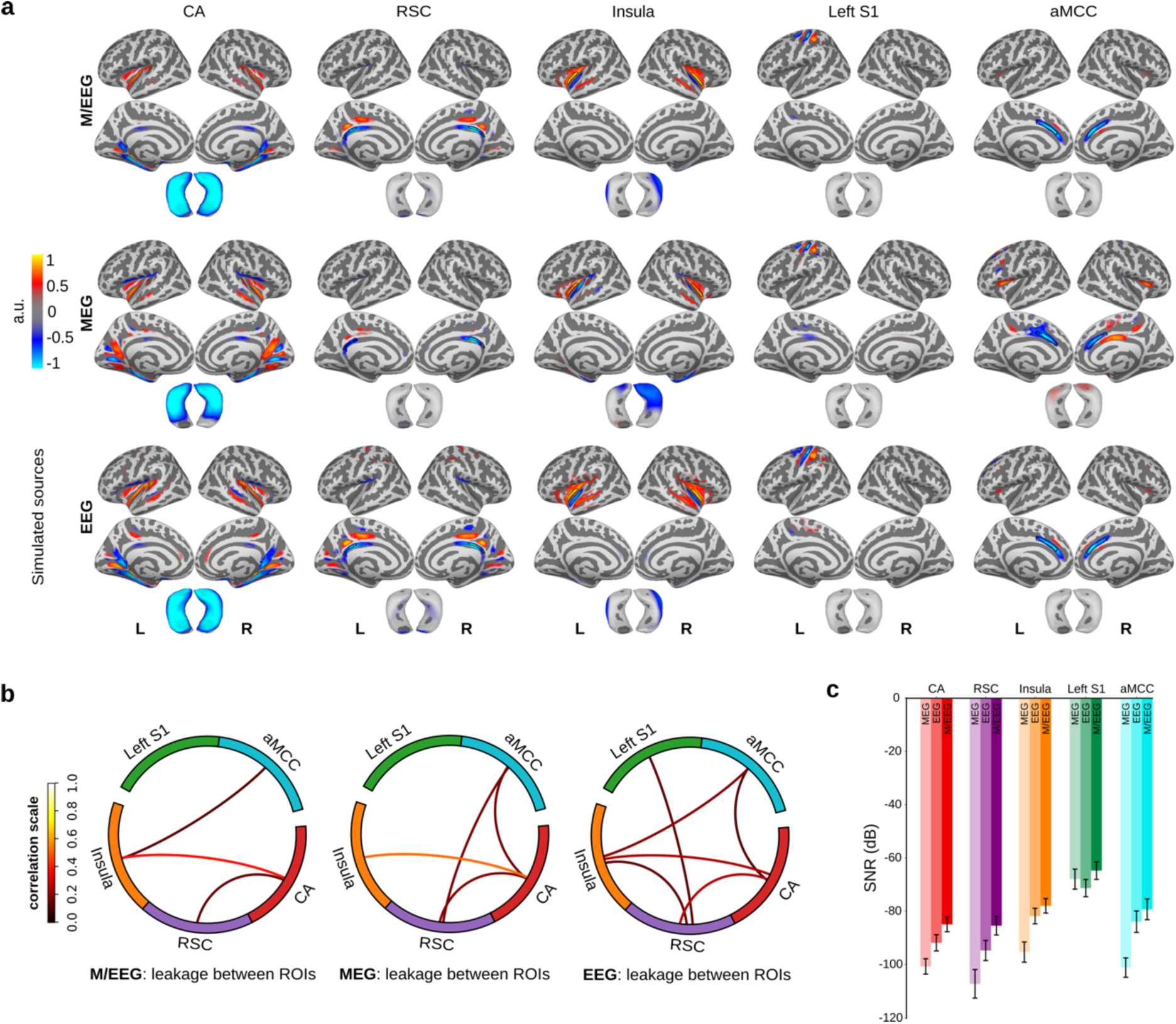
Relative spatial distribution of simulated source activity. (a) Dynamic statistical parametric maps (dSPM) for simulated M/EEG, MEG, and EEG (N=19). The maps are normalized to their peak amplitude in each ROI (i.e. the absolute cutoff is different for each single map and modality) to highlight spatial distribution rather than amplitude and signal-to-noise ratio. These maps represent simulated sources in the bilateral cornu ammonis (CA), bilateral retrosplenial cortex (RSC), bilateral insula, left primary somatosensory cortex (S1), and bilateral anterior midcingulate cortex (aMCC). (b) Point spread analysis for combined M/EEG and separate MEG and EEG, based on the simulations shown in a and illustrated with a circular graph, using a correlation cutoff value of 0.1. (c) Signal-to-noise (SNR) ratio for simulated activity over realistic noise within the ROI used for simulation, comparing EEG, MEG, and combined M/EEG.

Besides the interaction between sources regions, the SNR of source reconstruction also depends on the interaction with the ongoing M/EEG, including brain activity from many other sources. This was independently evaluated by calculating the SNR of simulated activity relative to real background M/EEG (Fig. 9c). While EEG generally provides a better SNR than MEG for midline sources and hippocampus (CA), the SNR is nevertheless enhanced in the combined M/EEG analysis compared to either method alone. The worst SNR in MEG among the five ROIs is observed for RSC, but still MEG contributes to enhance the combined M/EEG SNR in RSC. Conversely, the S1 source is the only one that shows better SNR for MEG than EEG, with relatively little SNR added by the combined M/EEG in comparison to the deep sources.

## Discussion

These results are consistent with the notion that the P3 is predominantly generated by a set of sources that can all be classified as part of the limbic system, including in particular RSC and hippocampus. Insula and aMCC may also be active in the P3 time window, but their contribution is somewhat later and negative-going in frontal mid-line electrodes. Below we will discuss the evidence for these sources and their consequences for the functional interpretation of the P3.

Based on the data of the present study, the RSC is the dominant source of the P3 in EEG. Further support for the RSC source comes from a recent study of tone-in-noise detection (Doll et al. 2024), such that a main P3 source in RSC has now been observed with auditory (Das et al. 2024) and visual oddball tasks, as well as with a near-threshold, single-tone detection task, providing a reproduction across different paradigms and modalities.

While previous experiments all used target stimuli, the detection of which was coupled to a button press, the present data add a distractor condition without button presses. While the amplitude of this response and, consequently, the signal-to-noise level of the source analysis, were much lower than that of the target-elicited P3, the results are nevertheless similar, suggesting a source in RSC for the distractor-elicited P3 in EEG as well.

The detailed comparison of EEG and MEG provided in the present study further shows that the sensitivity of MEG is very low for an RSC source, implying that other P3 sources are more dominant in MEG. The most prominent contributor to MEG in the P3 time-window was S1, which would not be considered a component of the P3, because we expect this component to depend on the button presses required for reporting and to be absent if the targets were e.g. counted instead. Unexpectedly, the sparse model fit suggests that S1 activity is also contained in the distractor P3, possibly related to actively suppressed motor responses in these no-go trials (Smith et al. 2008).

Including a hippocampal source space into the present analysis of P3 suggests a source with negative-going activity at the surface of regions CA1-4. The distributed source model used here additionally shows positive-going activity in the subiculum and presubiculum, whose orientation is opposite to CA 2-4, and then again negative-going activity in the convexity of parahippocampal gyrus. Such an alternating pattern is typical for a focal source in one side of a sulcus, and was similarly seen in simulations based on a source in CA. The hippocampus is generally a cortical structure (allocortex) with parallel aligned pyramidal cells (Palomero-Gallagher et al. 2020), and the extended source model used in this paper assumes that hippocampal sources can be modeled similar to cortical sources, which is well grounded on theoretical and experimental work for CA3 (Murakami et al. 2003). The anatomical modeling of the hippocampus in the source model introduced here allows for a more detailed comparison with predictions from intracranial recordings. In analogy to neocortex, the cortical surface of the CA is on the inside of the hippocampus formation.

The presence of P3-like activity in hippocampus is well supported by intracranial EEG (Halgren et al. 1980; Stapleton and Halgren 1987; McCarthy et al. 1989; Grunwald et al. 1999; Fell et al. 2005; Axmacher et al. 2010; Barbeau et al. 2017). These studies consistently reported negative-going P3 activity for depth electrodes placed along the center of the hippocampal formation, with a maximum in the posterior CA (Halgren et al. 1980; Stapleton and Halgren 1987; McCarthy et al. 1989; Grunwald et al. 1999; Ludowig et al. 2010; Barbeau et al. 2017). While these studies also reported contacts in the subiculum, or more generally in parahippocampal gyrus, no consistent activity was reported there, such that the hippocampal P3 source was expected to produce an open field. Only one study reported a second source in the anterior subiculum (Ludowig et al. 2010) with a negative-going polarity. This source could cause some cancelation with the hippocampus source because of its opposite orientation, but provided that the polarity in our source analysis was positive-going in the subiculum, it more likely reflects spread from the hippocampus rather than a genuine source in the subiculum.

The average dipole orientation (normal vector) of the hippocampus ROI is towards the frontal scalp (Fz; cf. Fig. 7a). Based on anatomical considerations (Iglesias et al., 2015), this hippocampal P3 source was expected to correspond more to the dorsal (rather than to the lateral) side of hippocampus, comprising mostly CA2-3 and some CA1 (cf. Fig. 2).

The finding that lesions of the hippocampus do not change the P3 in EEG (Johnson 1988; Onofrj et al. 1992) only pertains to the prominent component at Pz. The latter finding is well explained by our source model, because Pz activity appears to be mostly generated in RSC, whereas the peak of a simulated CA source is at Fz. Instead, the frontal maximum matches well with the EEG scalp distribution suggested for the P3a, which was demonstrated to be reduced by unilateral medial temporal lobe lesions (Knight, 1996). The present data thus support the conclusion of Knight that the hippocampus contributes to the P3 in EEG. This applied in particular to the “P3a” scalp distribution evoked by novel stimuli, but the lesion patients in that study also showed slightly reduced frontal activity for a target stimulus. Conversely, we suggest that the RSC is the main source of the parietal topography that is typically labelled as “P3b”. This topography remained unaffected by temporal lesions and was still observed for a novel stimulus when the frontal part was reduced (Knight, 1996). A problem for the definition of the P3a and P3b components (Squires et al. 1975; Polich 2007) is that it confounds two different scalp topographies (parietal versus frontal) with two different stimulus contexts (targets versus rare/novel distractors). However, it is obvious that the hippocampal, intracranial P3 has been directly recorded with target-detection paradigms (Halgren et al. 1980; Stapleton and Halgren 1987; McCarthy et al. 1989; Grunwald et al. 1999), which are used to obtain the P3b in scalp EEG (Polich 2007). Moreover, a hippocampal P3 has been similarly shown for auditory targets and distractors, even though it was not labeled P3a in the latter case (Halgren et al., 1995b). Based on the data presented in this paper, it appears that the RSC and hippocampal components of the P3 are present for the target and non-target P3 alike, but with a different amplitude dominance. Since the distractors used in the present experiment evoked a relatively small P3 and no frontal maximum, it cannot be used as example of a typical P3a. Nevertheless, it is conceivable that the hippocampal P3 could be more prominent in non-target and yet stronger in novelty paradigms (Courchesne et al., 1975; Knight, 1996), shifting the maximum from Pz to Cz, or even to Fz (Polich 2007). Such a shift of maximum can be easily demonstrated by adding simulated RSC and hippocampus activity with different relative amplitudes. A strong overlap of brain areas active for target and non-target stimuli has similarly been observed when target and non-target oddball stimuli were studied in fMRI (Kim 2014). Differences between rare target and non-target stimuli have nevertheless been observed, but are not inconsistent with the hypothesis outlined above, which relates specifically to the responses observed with maxima at Pz and Cz. Part of the sources described for novel stimuli have been observed in and around modality specific (auditory) cortex with fMRI and intracranial EEG (Sabri et al. 2006; Nourski et al. 2024), likely co-occuring with the P3a in EEG (Rinne et al. 2006). The P3 network suggested in this study for targets appears to be stable between the auditory and visual modalities, and is likely supramodal. Modality specific activity in the P3 latency range has been previously observed in auditory cortex for targets (Das et al. 2024) and non-targets (Kretzschmar and Gutschalk 2010) using MEG, as well, and is interpreted by us to be independent of the supramodal P3 activity. A visual equivalent may likely exist, but could not be identified in the present study. While activity in the fusiform gyrus was observed for visual targets in the P3 latency range, it was similarly observed for auditory targets and a simulated CA source, making it more likely to be spread from CA rather than focal visual activity.

There has been a long discussion on whether deep sources like the hippocampus can be recorded in MEG at all. The resolution to this question appears to be highly individual: Some subcortical sources like the auditory brainstem response can be recorded with MEG (Parkkonen et al. 2009; Dykstra et al. 2016) and are clearly identified not only by their source location but also by their early latency. A series of MEG studies had previously reported a medial-temporal-lobe source of the P3 based on dipole source analysis (Rogers et al. 1993; Tarkka et al. 1995; Basile et al. 1997), and more recently hippocampus activity was reported in MEG studies of hippocampal replay (Liu et al. 2019; Wimmer et al. 2020).

Further support for the hippocampus being accessible to MEG comes from methodological studies (Attal and Schwartz 2013; Krishnaswamy et al. 2017; Pizzo et al. 2019; Ruzich et al. 2019). Importantly, the detection of deep sources is not limited to magnetometers but similarly possible with (planar) gradiometers (Hunold et al. 2016). While MEG appears to be almost blind to the RSC source in comparison to EEG, which is in accordance with forward-model based sensitivity maps (Goldenholz et al. 2009), the situation is more complex for hippocampus. Available sensitivity estimates for the hippocampus have been calculated separately for radial and tangential source currents (Piastra et al. 2021): While the SNR is about 20dB lower in MEG than EEG for radial source currents, the disadvantage of MEG is much lower for tangential source currents. Based on a three-compartment model as used in the present study the SNR is about 6dB lower in MEG than EEG. When the CSF is additionally modeled, the disadvantage decreases to less than 3dB (Piastra et al. 2021). Based on the present data, we expect that the hippocampal source has strong tangential source currents and is therefore accessible to both, EEG and MEG. Using forward models that consider the CSF (Vorwerk et al. 2014) and individually calibrate the conductivity of the skull (Antonakakis et al. 2019; Antonakakis et al. 2020) may further improve the modelling of deep sources in particular for EEG but also MEG (Vorwerk et al. 2025), and may further improve the SNR that can be achieved by recording hippocampal activity with combined M/EEG.

Even with combined M/EEG, however, spread remains a problem for the interpretation of source analysis results. In the present study this is most critical for the spread observed between hippocampus and insula. Based on our model, the insular source has a positive-going polarity at the cortical surface, while the activity in RSC, CA, and aMCC is negativ-going. In case of CA, this is further supported by intracranial studies (Halgren et al. 1980; Stapleton and Halgren 1987; McCarthy et al. 1989; Grunwald et al. 1999). Biophysical models of evoked potentials suggest that cortico-cortical input into layer 1 dendrites of pyramidal cells is associated with negative-going activity at the cortical surface, whereas thalamo-cortical input into deeper layers is associated with positive-going activity (Jones et al. 2007). Therefore, positive-going activity in the insular P3 source would imply that the underlying mechanism differs in insula, possibly including thalamic input, whereas the negative going-polarity in RSC, aMCC, and - to the degree that the cortex model can be applied here - hippocampus would suggest the integration of incoming cortical information. Alternatively, the positive-going polarity in the insula ROI could be explained by spread from the CA source (Fig. 9a), but on the other hand, the peak latency was earlier in the CA compared to the insular P3 source, which argues for two independent sources. Moreover, intracranial recordings have confirmed insular activity in the P3 latency range (Citherlet et al. 2020), providing independent evidence for a potential insular source.

Beyond the question of P3 sources, which was the main focus of this paper and awaits confirmation from future work, another important question is what these sources indicate about the role in human cognition that the P3 indexes. Based on the peak latency of the sources, we found that hippocampus and RSC on the one hand, and insula and aMCC, on the other hand, have similar peak latencies. The first of these pairs, the RSC and hippocampus, have been shown to be densely and reciprocally connected (Kobayashi and Amaral 2003; Kobayashi and Amaral 2007). Traditionally, RSC and hippocampus have been thought to be jointly involved in episodic memory encoding and spatial navigation (Vann et al. 2009). The RSC is also thought to be generally involved in the coding of significant events (Smith et al. 2018), which could be easily linked with the presence of RSC and hippocampal P3 activity for both target and salient non-target events. The second pair, aMCC and insular cortex, are frequently coactivated during resting-state fMRI activity and are variably labeled as the ventral attention (Menon and Uddin 2010; Thomas Yeo et al. 2011), or cingulo-opercular/insular network (Sadaghiani et al. 2009; Doll et al. 2025 Mar 23). While RSC and hippocampal activity build up towards the button press and then decrease again, the maximum activity in aMCC showed a longer latency after the button press. The insular latency observed in the present visual study was longer than in RSC and hippocampus and numerically earlier, but not significantly different from aMCC. This is similarly evident when source time-courses were computed for targets, averaged by the button presses (Fig. S7b). Note, that insula activity was somewhat earlier in the previous auditory study (Das et al. 2024), suggesting overall that aMCC activity may succeed insular activity in latency. Moreover, the spread analysis demonstrated some interaction between insula and aMCC, which should be considered when interpreting the potential neural synchrony between these sources.

### Limitations

The source analysis techniques applied in the present study have inherent limitations because of the non-unique solution of the inverse problem in a volume conductor (Hämäläinen et al. 1993). While techniques that do not have such limitation, like fMRI and intracranial EEG, support the sources in RSC (Das et al. 2024; Doll et al. 2025 Mar 23), hippocampus (Halgren et al. 1980; Stapleton and Halgren 1987; McCarthy et al. 1989; Grunwald et al. 1999), and insula (Citherlet et al. 2020), we cannot exclude that further sources contribute to the P3. A number of such sources have been suggested based on fMRI and intracranial EEG (Halgren, Baudena, Clarke, Heit, Liégeois, et al. 1995; Halgren, Baudena, Clarke, Heit, Marinkovic, et al. 1995; Bledowski et al. 2004; Mulert et al. 2004), but were not confirmed in our source analysis based on M/EEG. Alternative source configurations can therefore not be ruled out at this point. For example, it is possible that the contribution of other sources is more variable across participants and is therefore not observed in the group-level analysis used in the present study (Hietala et al. 2024). Also, the spatial precision of M/EEG is limited, such that the exact location of the sources suggested here should be interpreted cautiously and considering that M/EEG cannot estimate the extend of a source. Finally, the analysis of the distractor P3 in the present study was exploratory and will require confirmatory studies to probe the source configuration for the P3a and P3b variants proposed above.

## Supporting information

Suppl. Results

## Author contributions

Diptyajit Das (Conceptualization, Experimental design, Data curation, Investigation, Methodology, Formal analysis, software, Visualization, Writing - original draft & editing), Matti S. Hämäläinen (Methodology, Validation, Writing - reviewing & editing), Andrew R. Dykstra (Validation, Writing - reviewing & editing), Andre Rupp (Validation, Writing - reviewing & editing), Alexander Gutschalk (Conceptualization, Experimental design, Methodology, Validation, Writing - original draft & editing, Funding acquisition, Supervision).

## Data availability

The raw and processed MEG and EEG data and scripts will be shared upon publication and can be accessed on heiDATA, the open research data repository of Heidelberg University, Germany.

## Abbreviations

aMCC: anterior midcingulate cortex
CA: cornu ammonis
dSPM: dynamic statistical parametric mapping
EEG: electroencephalography
MEG: magnetoencephalography
MNE: minimum-norm estimate
ROI: regions of interest
RSC: retro-splenial cortex
S1: primary somatosensory cortex.

