## Supplementary material for "Combined MEG and EEG suggest a limbic source network of the P3 including retrosplenial cortex and hippocampus": Suppl. Results

### Supplementary data:

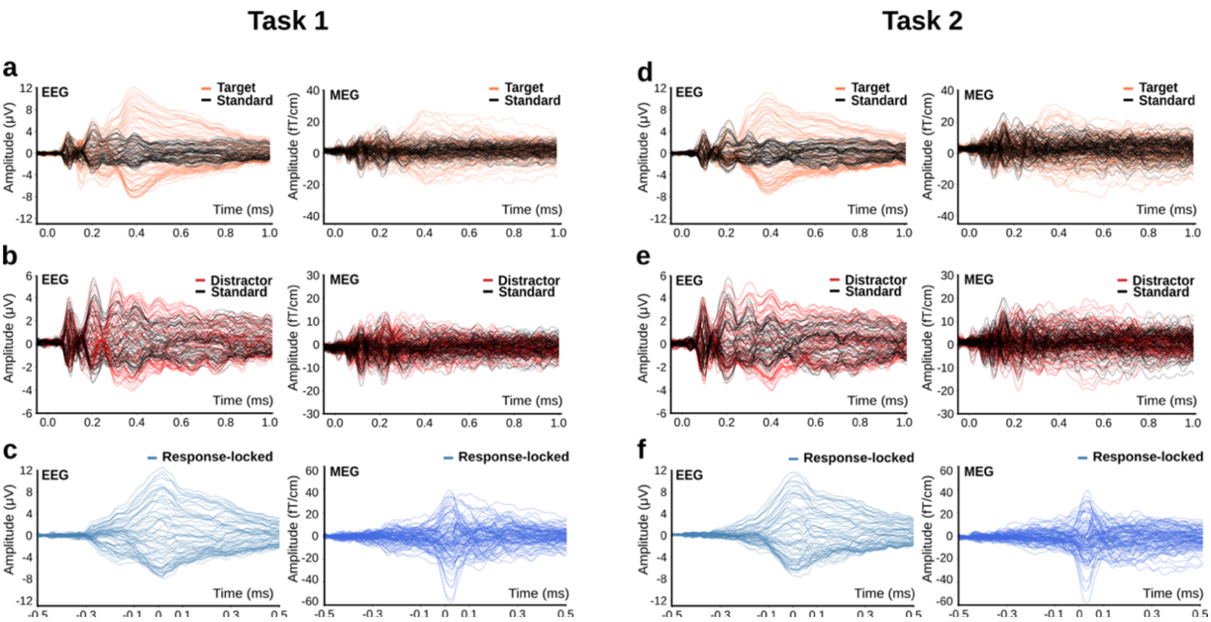


**Fig. S1.** Comparison of the evoked signals for paradigm variants. Variant 1 (left; a, b, c) and Variant 2 (right; d, e, f) show highly similar brain responses for targets (coral; a, d), standards (black), distractors (red; b, e), and stimulus-locked evoked signals (blue; c, f). All data in the main paper represent a combination of the two variants (cf. Figure 2).

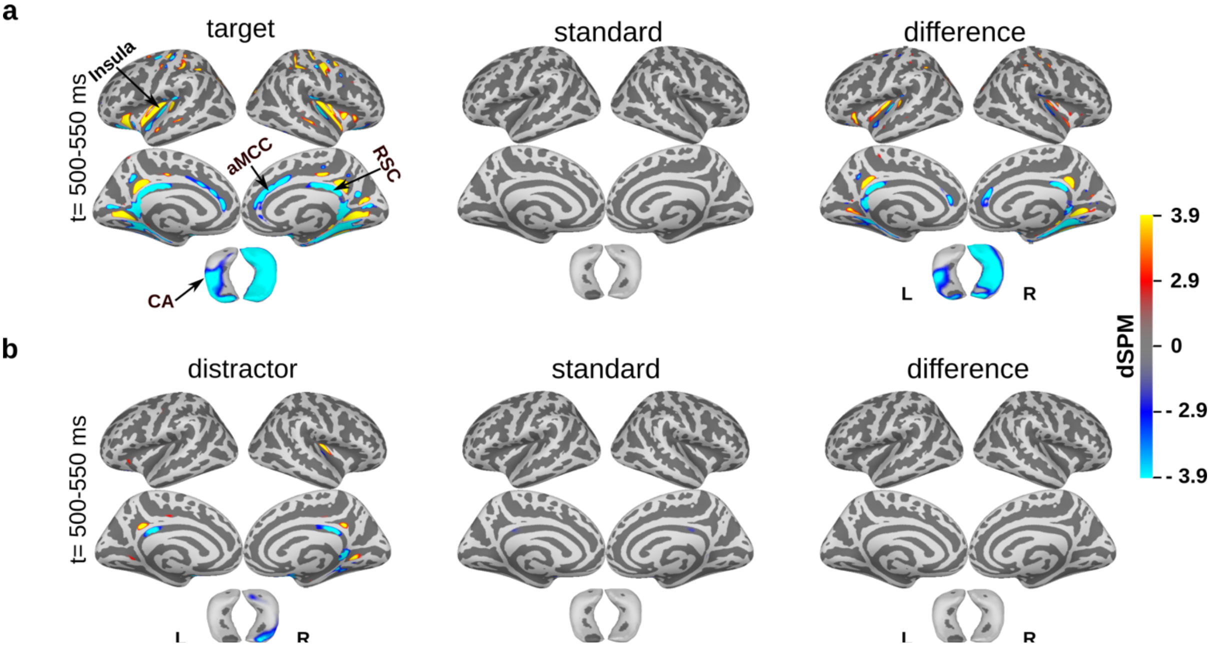


**Fig. S2.** Grand-average dSPM maps in a later time window (500 – 550 ms). The later time window was chosen to test if activity in anterior mid-cingulate cortex (aMCC) was observed with the visual oddball paradigm, similarly as in the previous auditory study. (a) Combined M/EEG dSPM maps ($p<0.01$; N=19) for targets (left), standards (middle), and the contrast between targets minus standards (right) in the time window 500–550 ms after stimulus onset. Besides showing activity in anterior mid-cingulate cortex (aMCC), this analysis also shows somewhat stronger insular activity for target stimuli than seen at the P3-peak latency in Figure 4. (b) Same arrangement as in a for distractor stimuli.

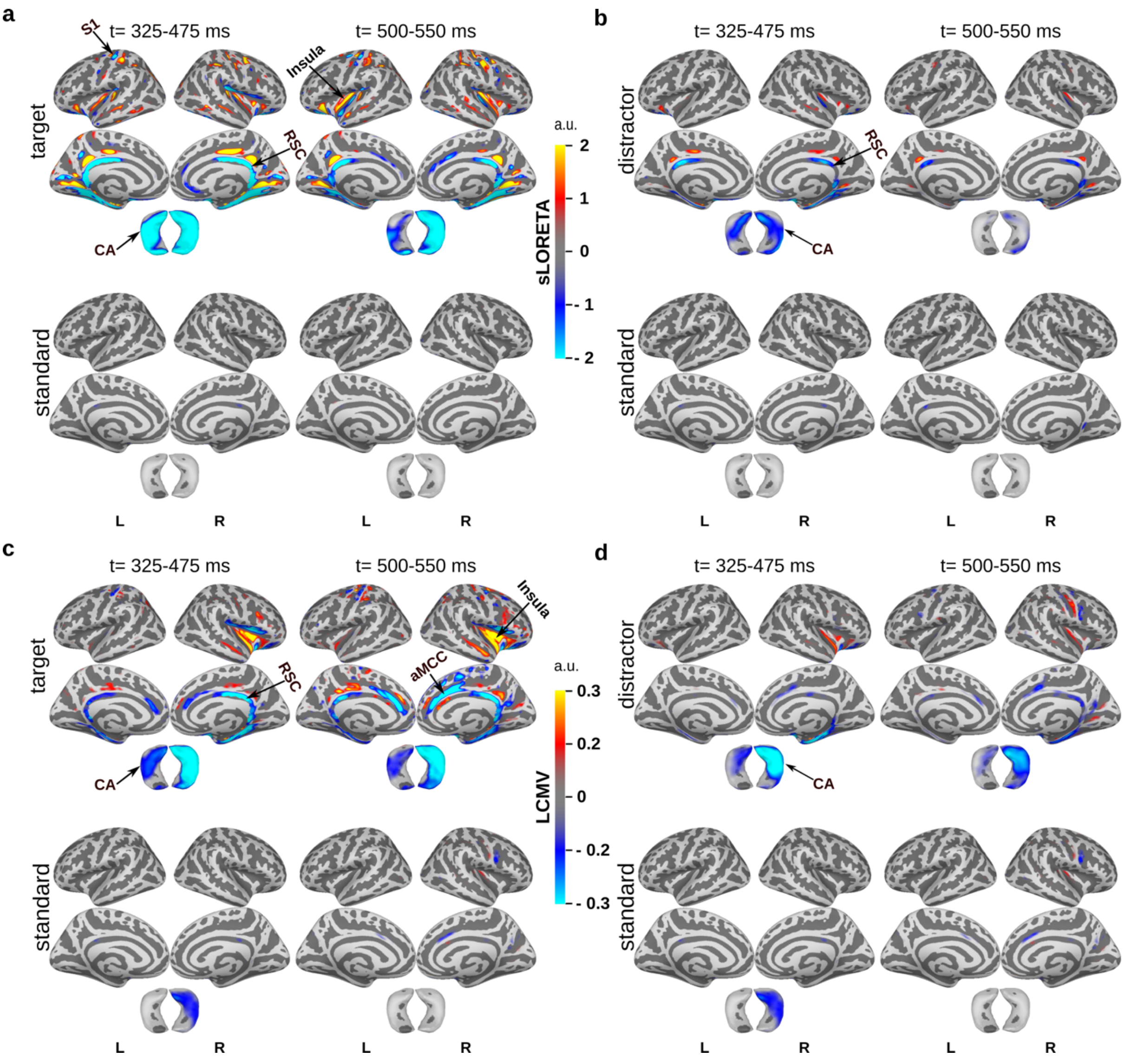


**Fig. S3.** Alternative source analysis methods (N=19). These group-activation maps are similar as the dSPM maps in Figure 3 and S2, but with alternative source analysis techniques. The maps are based on combined M/EEG in the time window 325 - 475 ms and 500 - 550 ms. Source maps are shown for target and standard (a, c), and for distractor and standards (b, d). (a, b) Analysis using a Borgiotti-Kaplan beamformer (LCMV; Sekihara and Nagarajan, 2008). (c, d) Analysis using sLORETA (standardized low-resolution brain electromagentic tomography; Pascual-Marqui, 2002). Note that, although both methods produce noise-normalized z-scores, the numerical values cannot be directly compared between methods and with the dSPM maps in Figures 3 and S2.

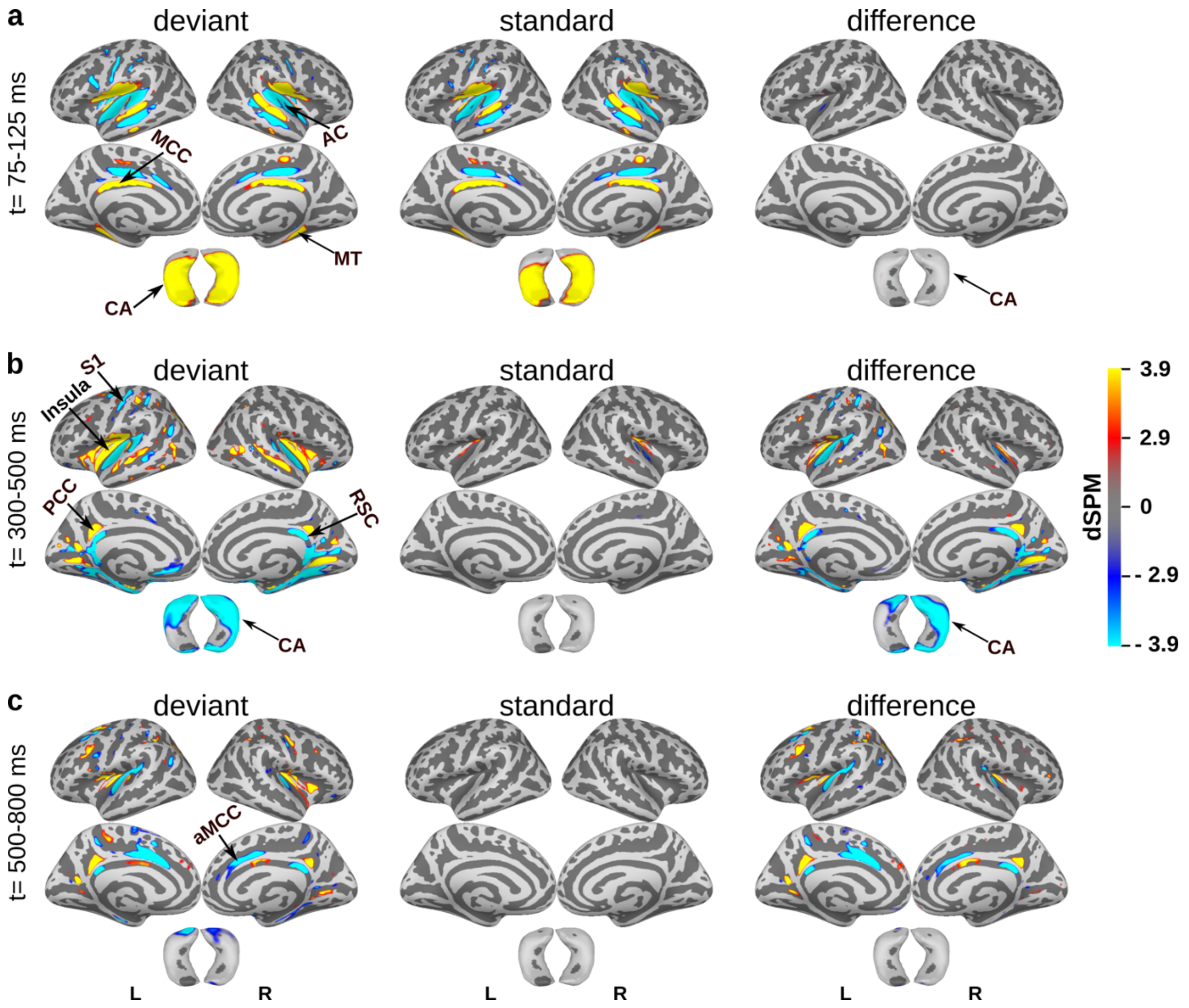


**Fig. S4.** Reanalysis of the data from Das et al 2024 (N=12) with the extended source space including the hippocampus (taken from Das, 2025). (a) The time interval 75 - 125 ms includes the auditory-evoked N1 response, which is known to be generated in auditory cortex (AC). This response was not significantly different between standards and deviants. As can be seen in the maps, this response also spreads to the CA regions of the hippocampus, but with a positive-going polarity. This is because of the similar orientation (b) The 300 - 500 ms time interval includes the P3 peak at Pz across participants and shows the previously reported activity in retro-splenial cortex (RSC) and insula. Additionally, activity in the anterior cornu ammonis (CA) is observed in the hippocampus source space. (c) The subsequent time interval 500 - 800 ms shows additional activity in the anterior mid-cingulate cortex (aMCC), whereas activity in the other sources fades away.

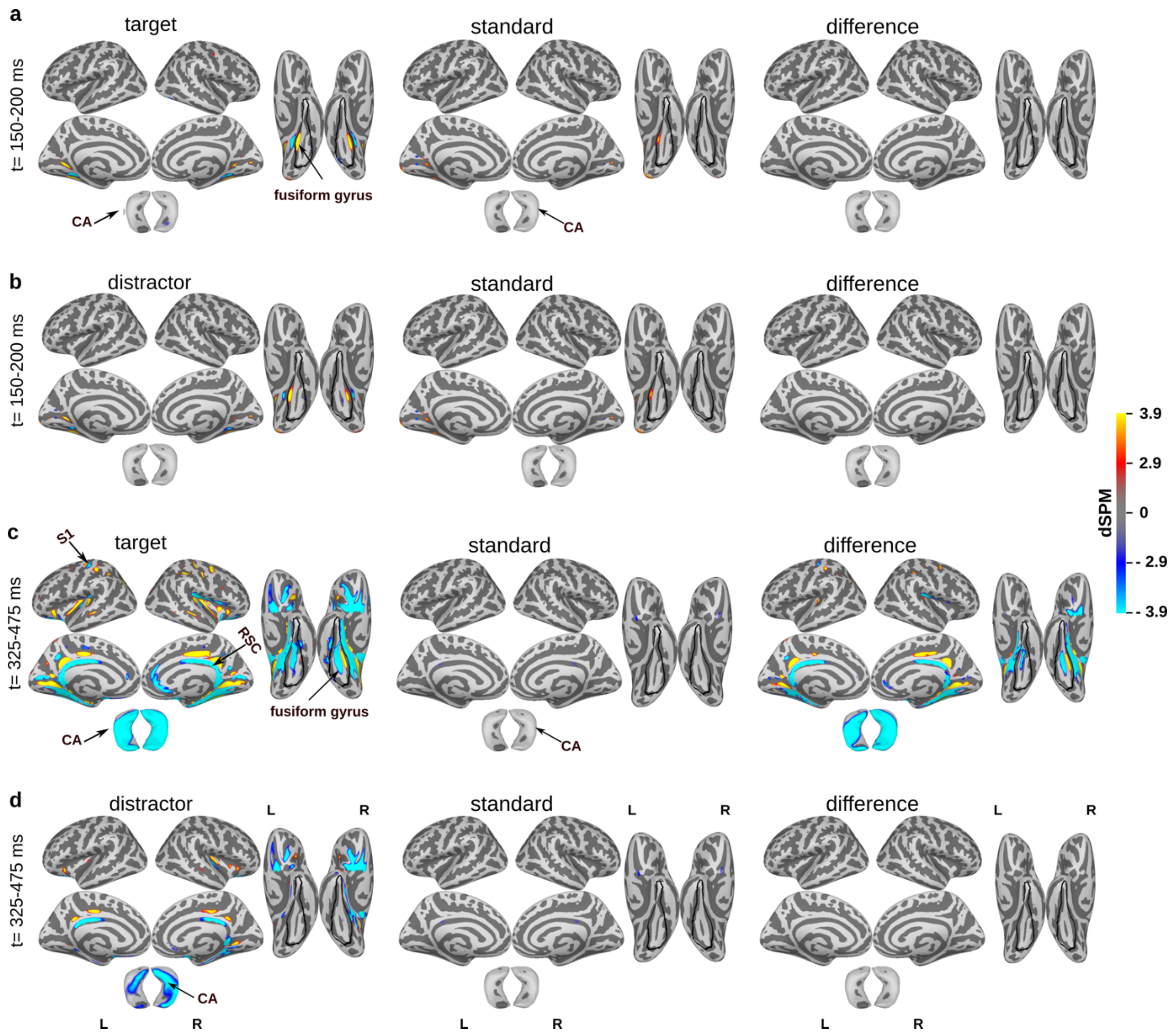


**Fig. S5.** Grand-average dSPM maps with an earlier time window and a view on the ventral temporal lobe added to evaluate activity in fusiform gyrus and lingual cortex. Combined M/EEG dSPM maps (p<0.01; N=19). The time window 150 - 200 ms subsequent to the P100 was chosen to focus on sensory activity in the fusiform and lingual gyrus. Mapping of the P100 did not reveal significant activity at this cutoff. Left column: (a, c) targets; (b, d) distractors. Middle column standards. Right column: (a, c) target - standard; (b, d) distractor - standard. (a, b) Sensory time window (150 - 200 ms); (c, d) P3 time window (325 - 475 ms).

(a, b) in the sensory time window, focal activity is observed at the border of lingual cortex and fusiform gyrus, and less prominent in the calcarine sulcus. No significant spread to CA is observed at the cutoff used (but some spread to posterior CA is observed when the threshold is lowered). The activity is similar for targets and distractors. Activity evoked by standards is weaker, but the difference is not significant.

(c, d) maps P3 with identical settings as in Fig. 4 with the ventral view of the temporal lobe added. As can be seen there, negative going activity is also observed in the fusiform gyrus with a similar pattern as for the sensory activity but more wide spread. This activity may as well represent spread from CA, as is suggested by simulations (not shown). It cannot be excluded that this activity represents real visual processing, but typically activity in this region (V4) is stronger in the earlier time window. Based on (a, b) it is therefore unlikely that CA activity receives major spread from fusiform gyrus, but this cannot be certainly distinguished on the source analysis in the P3 time window.

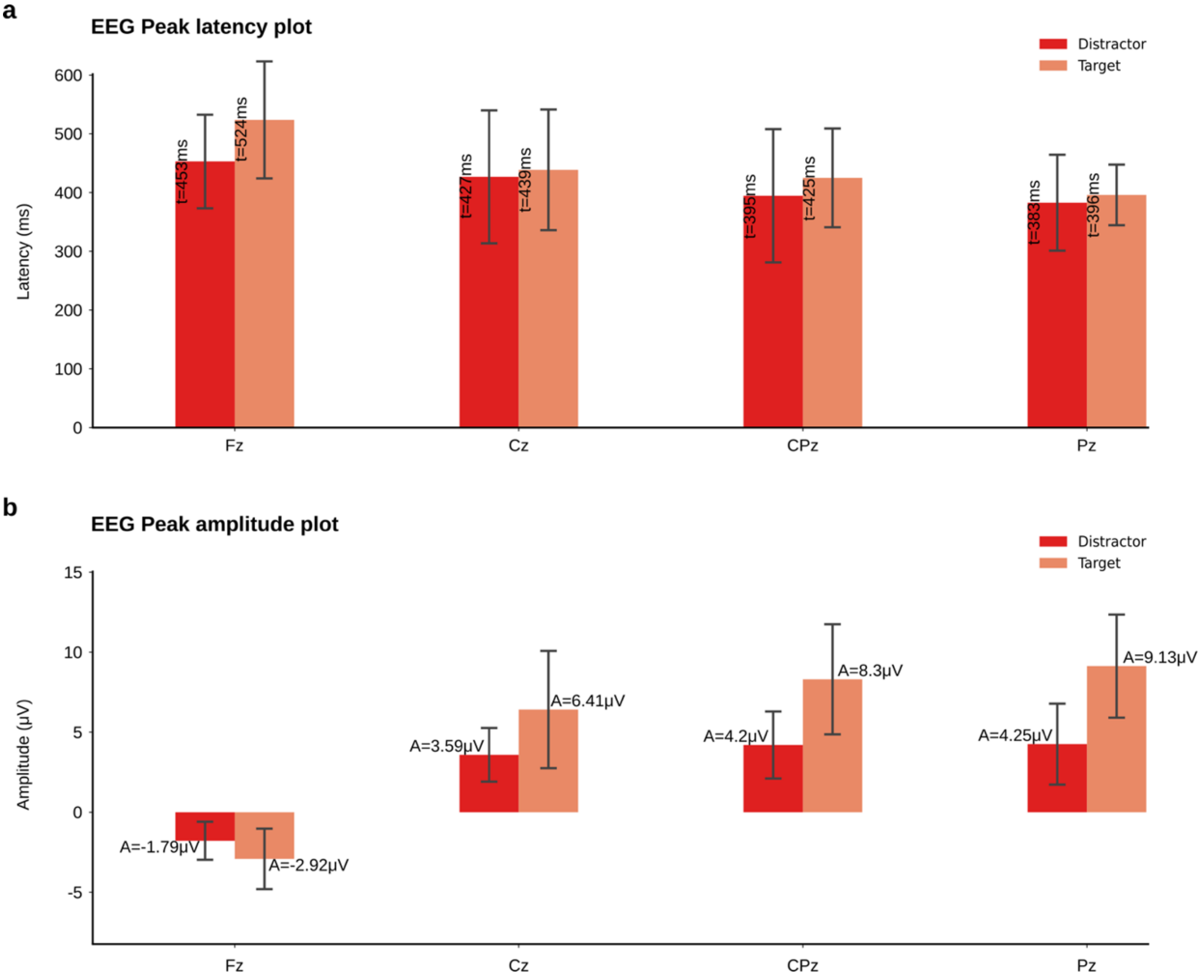


**Fig. S6.** P3 peak latencies and amplitudes in midline EEG electrode positions (Fz, Cz, Cpz, and Pz). (a) Peak latency and (b) peak amplitude (mean ± standard deviation; N=19) of the P3 measured in the midline sensors (Fz, Cz, Cpz, and Pz) for targets (coral) and distractors (red). Targets elicit a stronger P3 than distractors (b), while the latency of distractors brain responses appears to be slightly earlier (a) in all midline sensors.

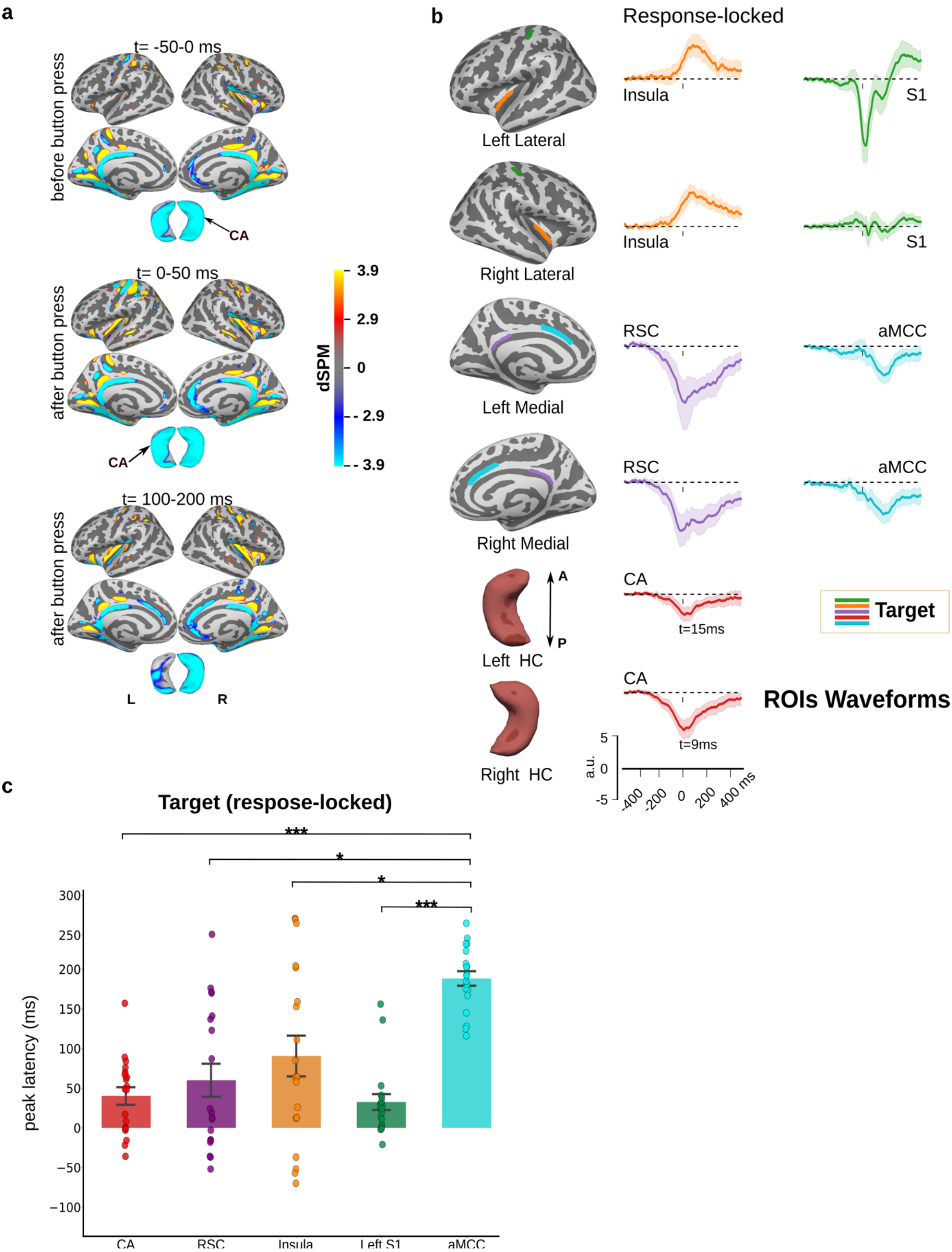


**Fig. S7.** Response-locked brain activity (N=19). (a) Combined M/EEG dSPM maps ($p<0.01$) for targets, averaged by the button press in the time window of 50 ms pre-button press (top) and 50 ms post-button press (bottom). (b) M/EEG source waveforms extracted from dSPM estimates across participants for response-locked targets. The ROIs (region of interest) are plotted in the inflated fsaverage brain atlas (right column). The same color code is used to represent each ROI and its corresponding source waveform. The ROIs include the insula, primary somatosensory cortex (S1), retrosplenial cortex (RSC), anterior midcingulate cortex (aMCC), and cornu ammonis (CA). The shaded areas represent the 95% confidence intervals, calculated using non-parametric bootstrap method. (c) Peak-latency analysis of ROI-based source time courses (mean ± standard error) for the response-locked activity. The same ROIs are included here as in (b). Significant differences between peak latencies for ROIs are indicated by asterisks (* $p < 0.05$, ** $p < 0.01$, *** $p < 0.001$) based on t-tests for repeated measures with Bonferroni correction for multiple comparisons.

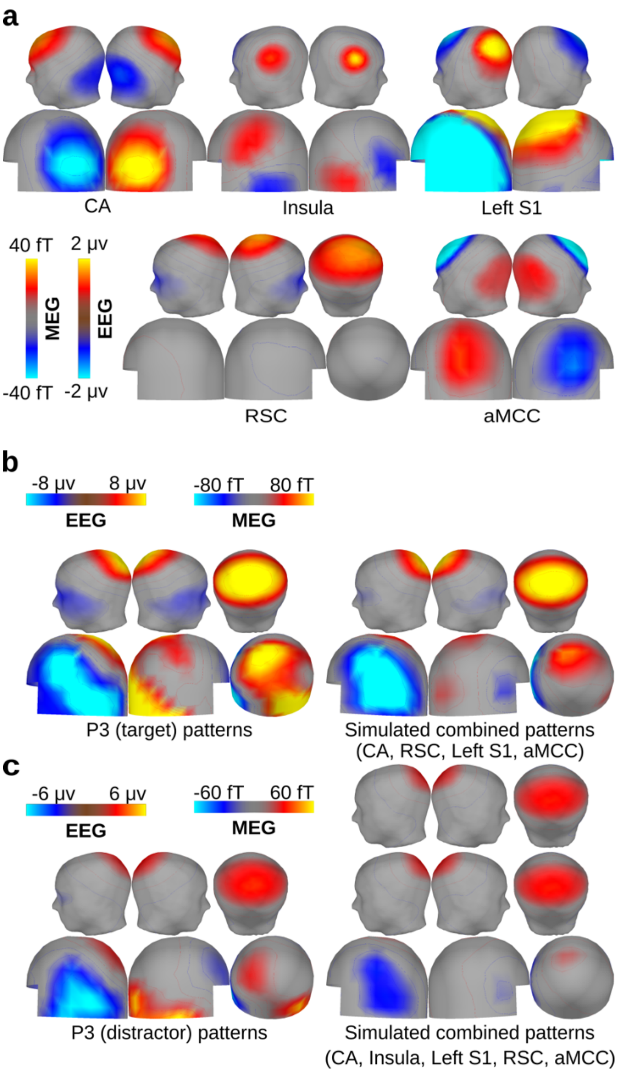


**Fig. S8.** Simulated M/EEG sources and sparse P3 source model, variant with a different head model. In this variant of the analysis, the skull conductivity was chosen to be 0.003, to compensate for an overestimation of the EEG amplitude when the CSF is not modeled. (a) Spatial pattern of scalp EEG (top) and virtual magnetometer maps (bottom) for MEG, derived from simulated sources (grand average, N=19) in the bilateral cornu ammonis (CA), insula, left primary somatosensory cortex (S1), bilateral retrosplenial cortex (RSC), and anterior midcingulate cortex (aMCC). (b) Grand-average (N=19) scalp EEG and virtual MEG magnetometer maps at the peak latency for the P3 in Pz for targets (left). Weighted (cf. table 1) combination of simulated sources in CA, insula, S1, RSC, and aMCC that best explain the P3 for targets (right). (c) Same arrangement as in b for distractor-evoked P3.

**Table S1:** Variant of the sparse model based on a head model with skull conductivity 0.003 (cf. Fig. S7). Relative weights and residual variance for the sparse model of the grand-average M/EEG for target P3 and distractor P3 with simulated, ROI-based sources. CA, cornu ammonis; RSC, retro-splenial cortex; S1, primary somatosensory cortex; aMCC, anterior middle cingulate cortex; EEG, electroencephalography; MEG, magnetoencephalography.

| **Condition** | **Simulated source activation strenght (nAm)** | | | | | **Residual variance (%)** | |
|---|---|---|---|---|---|---|---|
| | **CA** | **RSC** | **Insula** | **left S1** | **aMCC** | **EEG** | **MEG** |
| Target P3 | 32.0 | 105.5 | 0.0 | 18.0 | 16.2 | 5.7 | 33.9 |
| | 32.0 | 105.5 | - | 18.0 | 16.2 | 5.7 | 33.9 |
| | 18.8 | 110.2 | 14.8 | 18.8 | - | 5.3 | 35.2 |
| | - | 114.8 | 13.2 | 19.2 | 0.0 | 6.9 | 40.5 |
| | 52.0 | - | 0.0 | 15.2 | 28.7 | 76.1 | 35.7 |
| | 48.5 | 91.0 | 0.0 | - | 27.0 | 11.6 | 99.9 |
| Distractor P3 | 14.2 | 48.2 | 2.8 | 6.8 | 11.2 | 7.1 | 41.4 |
| | 16.0 | 47.2 | - | 6.5 | 13.2 | 7.1 | 41.4 |
| | 3.8 | 52.2 | 13.5 | 7.5 | - | 7.4 | 44.4 |
| | - | 53.2 | 12.8 | 7.2 | 0.0 | 6.5 | 49.5 |
| | 33.2 | - | 0.0 | 5.8 | 23.8 | 74.7 | 43.6 |
| | 20.8 | 45.5 | 0.0 | - | 18.5 | 8.6 | 93.8 |

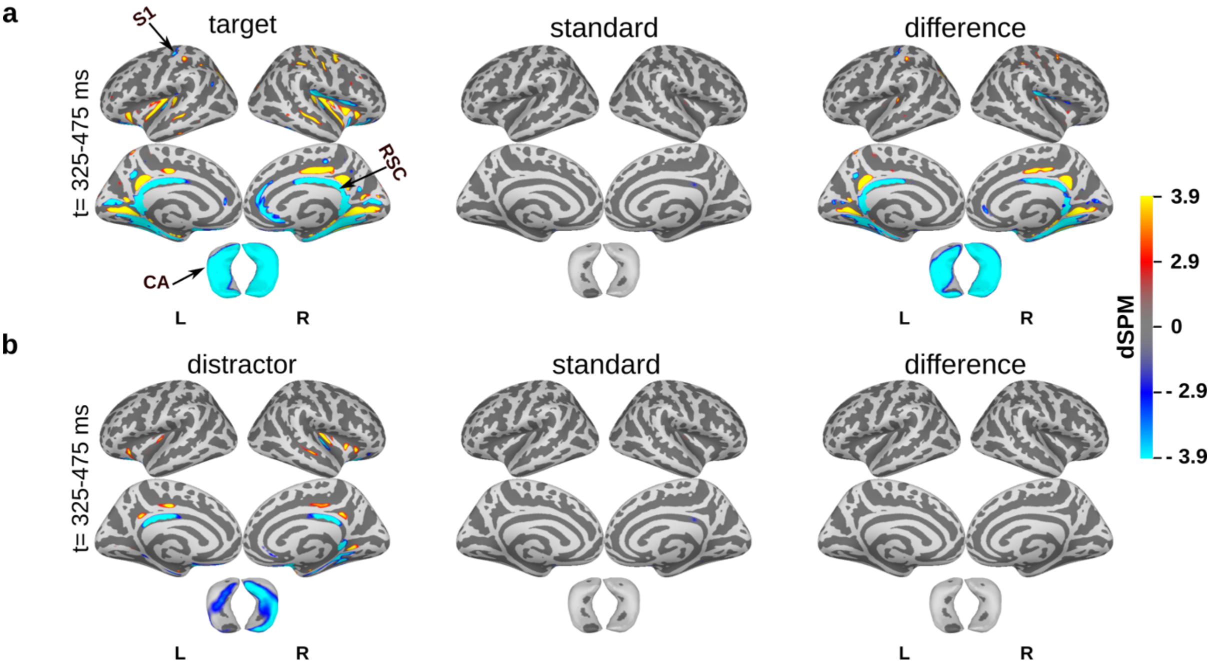


**Fig. S9.** Alternative dSPM analysis with a modified head model (skull conductivity 0.003). As can be seen in comparison to Fig. 4, the dSPM maps are quite robust to modifications of skull conductivity.

Grand-average dSPM activation maps for combined M/EEG (N=19). (a) Combined M/EEG dSPM maps (p<0.01) for targets (left), standards (middle), and the contrast between targets minus standards (right) in the time window 325–475 ms after stimulus onset. (b) dSPM maps for distractor stimuli, similarly arranged. While activity in RSC, insula, and CA is observed for distractors and not for standards, no significant difference is mapped between these two conditions (right column). The number of trials were equalized between targets, distractors, and standards prior to source estimation. Highlighted brain regions include retrosplenial cortex (RSC), primary somatosensory cortex (S1), insula, and the cornu ammonis (CA). Note that only the dorsal view of hippocampus is shown and that for technical reasons the activity is plotted on the outside, while the polarity shown corresponds to the cortical surface in the inside of hippocampus (cf. Fig. 2).

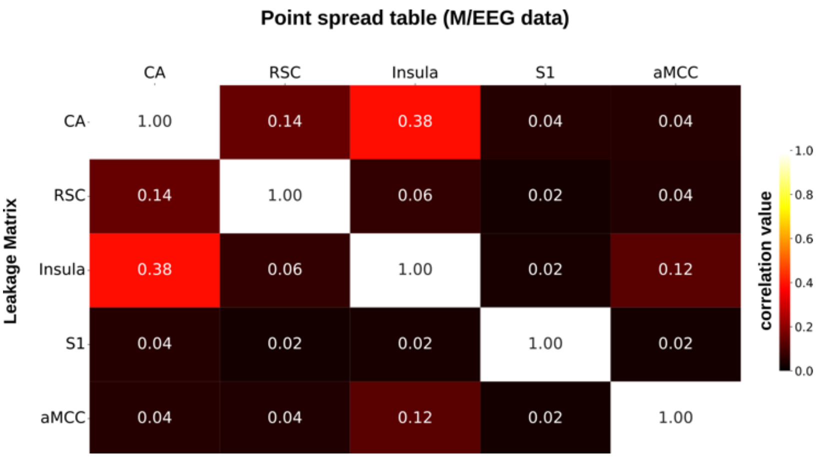


**Fig. S10.** Point spread analysis for dSPM between ROIs for combined M/EEG data. The matrix represents how the point source activity from one ROI (region of interest) would spread to other ROIs. The ROIs include the bilateral cornu ammonis (CA), retro-splenial cortex (RSC), insula, left somatosensory hand area (S1), and bilateral anterior midcingulate cortex (aMCC). The correlation values range from 0 to 1, with 1 indicating maximum leakage and 0 indicating no leakage.

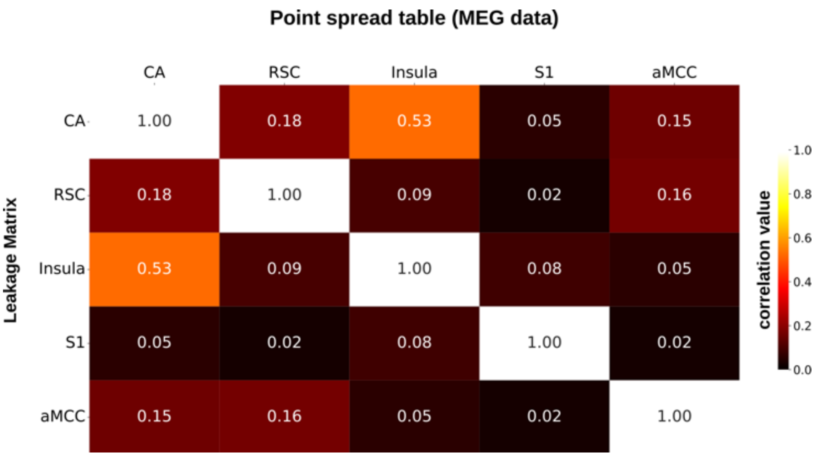


**Fig. S11.** Point spread analysis for dSPM between ROIs for MEG data only.

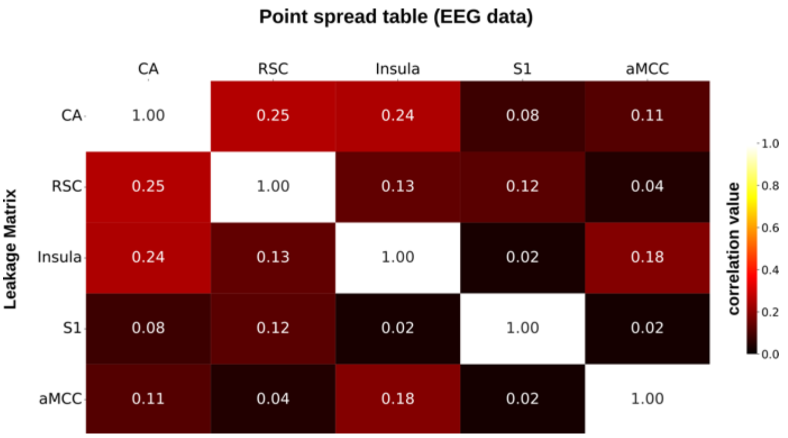


**Fig. S12.** Point spread analysis for dSPM between ROIs for EEG data only.
